# Remyelination therapy restores suppressed neuronal activity in a mouse model of demyelination

**DOI:** 10.1101/2025.02.03.636344

**Authors:** Aniruddha Das, Julie Borovicka, Hale Tobin, Rajaram Kshetri, Shruti Bhatia, Jacob Icardi, Ghazaal Tahmasebi, Priyanka Agochiya, Shriya Singh, Jordon Manworren, Philopater Isak, Willow Lev, Bruce D Trapp, Tara M DeSilva, Hod Dana

**Author notes:** These authors contributed equally to this work (Aniruddha Das, Julie Borovicka). Correspondence to: Hod Dana, Department of Neurosciences, Cleveland Clinic Research, Cleveland Clinic Foundation, Cleveland, OH 44195, USA.

## Abstract

Central nervous system demyelination contributes to disability in disorders such as multiple sclerosis, yet how myelin loss and repair alter neuronal function over time remains poorly understood. We longitudinally monitored hippocampal myelination and CA1 pyramidal neuron activity in the same mice for more than 100 days, spanning cuprizone-induced demyelination and subsequent treatment with clemastine, a clinically investigated remyelinating agent. Imaging-detectable myelin loss emerged only after several weeks, whereas neuronal activity declined within days, indicating that circuit dysfunction precedes myelin degeneration. Neuronal activity also exhibited sex-dependent biphasic trajectory, with greater early deficits in females and later deficits in males. Clemastine promoted axonal myelination and restored neuronal activity to pre-cuprizone levels. Longitudinal tracking of individual neurons across baseline, demyelination, and treatment further showed that recovery depended on cellular baseline properties: more active neurons exhibited larger declines during vehicle treatment and larger increases during clemastine treatment. Together, these findings identify sex-dependent neuronal responses to demyelination, link structural myelin repair with functional circuit recovery, and highlight heterogeneity in neuronal recovery associated with baseline activity, supporting the use of functional outcomes to evaluate candidate remyelination therapies.

## Introduction

Myelin is a multi-layered sheath produced by oligodendrocytes that insulates axons and enables the rapid propagation of action potentials (APs)^1^. In demyelinating diseases such as multiple sclerosis (MS), myelin degeneration leads to severe motor, visual, and speech impairments, as well as long-term cognitive decline^2,3^. Cognitive and memory deficits in MS are closely linked to hippocampal demyelination and atrophy^2,4^. While MS is recognized as a disease with a major autoimmune inflammatory component, basic experimental studies, human biopsy samples, and autopsy tissue indicate that primary oligodendrocyte dystrophy also contributes to disease pathogenesis^5,6^.

Rigorous study of demyelination-associated pathology requires appropriate animal models, particularly because human tissue is usually available only at advanced disease stages^7^. One widely used model is the mouse cuprizone model of demyelination^8^. Cuprizone, a copper chelator, induces selective death of mature oligodendrocytes in the brain, resulting in extensive demyelination in multiple regions, including the hippocampus^8–10^. Importantly, this model also allows investigation of remyelination, which occurs spontaneously after cuprizone withdrawal^8,11^. Previous studies have described gradual, brain-wide demyelination that typically emerges 2–3 weeks after cuprizone initiation and progresses for as long as cuprizone is administered^10^.

Cuprizone-induced hippocampal demyelination has been shown to alter CA1 neuronal function and impair long-term potentiation in *ex vivo* tissue slices^4,12^. Consistent with these findings, we previously demonstrated a reduction in hippocampal neuronal firing *in vivo* during cuprizone exposure, with partial recovery after return to a regular diet^13^. In the cortex, even incomplete remyelination driven by therapeutically enhanced oligodendrogenesis can be sufficient to restore functional responses^14^. However, a detailed, longitudinal understanding of how changes in neuronal myelination relate to neuronal activity in the same animal, on the time scale of days to weeks, remains lacking.

In this study, we longitudinally imaged CA1 pyramidal neurons *in vivo* using two-photon laser scanning microscopy (TPLSM) in transgenic mice expressing the red genetically encoded calcium indicator (GECI) jRGECO1a^15,16^, enabling us to monitor activity patterns with single-cell resolution across weeks. In the same animals, we simultaneously recorded third-harmonic generation (THG) signals, which provide label-free contrast from lipid-rich structures such as myelin and thus report myelination status with micrometer-scale resolution^17–19^. We show a rapid decline in neuronal firing rates (FRs) as early as 4 days after cuprizone administration, which precedes delayed and gradual hippocampal demyelination. This decline follows a sex-specific, biphasic pattern, more pronounced in females early and in males later in the cuprizone period. Whole-cell electrophysiological recordings demonstrated that acute cuprizone exposure does not directly modify neuronal excitability, suggesting that the observed changes were mediated by cuprizone-mediated effects on oligodendrocytes. Following cuprizone cessation, mice were treated with clemastine, an oral antihistamine that is being repurposed as a remyelinating therapy and is currently under evaluation in MS clinical trials^20^. Clemastine treatment promoted remyelination as well as restored neuronal FRs to pre-cuprizone levels. Our findings therefore identify sex-specific disruption of CA1 neuronal function as an early pathology that precedes demyelination and demonstrate that neuronal functionality can be protected by remyelination-enhancing treatment.

## Results

### Longitudinal monitoring of neuronal activity and myelin status

TPLSM is widely used to record activity from neurons expressing GECIs with single-cell resolution^21^. THG is a less commonly used nonlinear imaging method that provides complementary, label-free contrast from myelin-rich structures in brain tissue^17^. Here, we combined these two optical approaches to concurrently record firing of CA1 pyramidal neurons and monitor hippocampal myelination status in the same mice, thus enabling direct structure–function coupling over time (Fig. 1A-B). All mice were randomly assigned to two groups. The cuprizone group received a 0.3% cuprizone diet for 55 days, after which mice were randomly assigned to one of two subgroups that received 45 days of daily treatment with either the remyelinating drug clemastine or vehicle. The control group was maintained on regular diet for the entire study period and was divided into two subgroups, which received either clemastine or no treatment. We imaged mice before and during cuprizone consumption, and during treatment with clemastine or vehicle. All recording sessions included identical TPLSM and THG acquisitions and were conducted at defined timepoints to monitor longitudinal changes in neuronal activity, myelin status, and body weight (Fig. 1C, Fig. S1).

**Figure 1:**
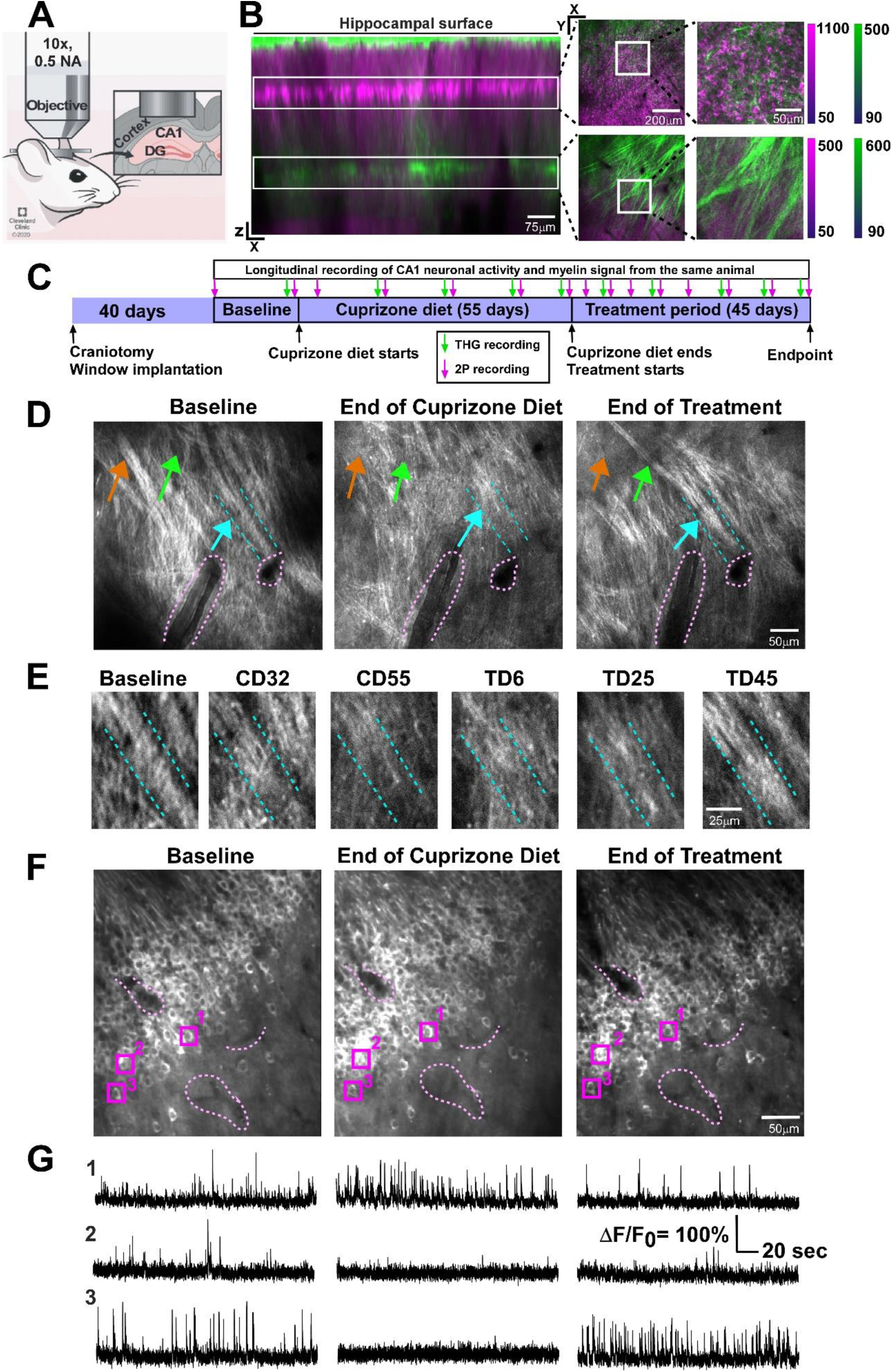
Longitudinal monitoring of hippocampal CA1 activity and myelination levels in a mouse model of de/remyelination. **(A)** Illustration of hippocampal window implantation to allow long-term access to the hippocampus of a transgenic Thy1-jRGECO1a mouse. **(B)** *In vivo* side view of a Thy1-jRGECO1a hippocampus shows CA1 neurons, axons, and dendrites expressing the red calcium sensor jRGECO1a (acquired by TPLSM and shown in magenta) and hippocampal myelin (acquired by THG and shown in green). Insets: horizontal views of the CA1 and myelinated fiber layers. **(C)** Timeline for neuronal activity and myelin recording sessions (indicated by magenta and green arrows, respectively). **(D)** Example large field of view THG images of the same hippocampal fibers illustrate the disappearance (Cuprizone Day 55, orange and cyan arrows), reappearance (cyan arrows), and *de novo* appearance of myelin fibers (green arrows) during the cuprizone administration and subsequent treatment periods. All images are contrast-enhanced for visibility; purple dashed lines indicate blood vessels used to reliably re-identify the FOV. **(E)** Example high-magnification THG images of the same hippocampal fibers shown in **D** illustrate the loss and recovery pattern of myelin during the cuprizone administration and subsequent treatment periods. **(F)** Example TPLSM images from the same FOV of CA1 neurons expressing jRGECO1a. The same cells are highlighted by the numbered magenta squares in the different images. Purple dashed lines indicate blood vessels used to reliably re-identify the FOV. **(G)** fluorescence traces from the cells shown in E. Action potential firing produces increases in the fluorescence signal, with pronounced differences in activity across the various stages of cuprizone consumption and treatment.

Myelin signal was quantified by counting pixels containing myelin and expressing their fraction of the total image pixels (myelin coverage), as well as by measuring the thickness and signal amplitude of individual myelinated fibers across recording sessions. This longitudinal monitoring revealed distinct patterns of myelin loss and recovery, including disappearance, reappearance, and *de novo* appearance of myelin fibers in the same field of view (FOV; Fig. 1D-E). Notably, the THG signal did not typically vanish entirely as demyelination progressed, but a weaker background signal was still apparent even after individual fibers were no longer detectable. TPLSM recordings were analyzed by measuring changes in GECI fluorescence and fitting a maximum-likelihood mode^22^ to infer the timing and number of action potentials in single neurons^13^. Consistent with previous reports in the mouse cuprizone model, we did not detect neuronal loss in our recordings^23–26^. Nonetheless, we observed altered cellular activity patterns during the cuprizone consumption and treatment phases (Fig. 1F-G; see Methods).

### Sex-specific patterns of CA1 activity suppression during cuprizone consumption

Following cuprizone consumption, myelin signal coverage gradually declined from baseline, becoming significantly reduced after 32–46 days on the diet; male mice showed lower myelin coverage than females from cuprizone day 32 (CD32) onward, but the difference was not significant (Fig. 2A). At the level of individual myelinated fibers, signal amplitude decreased and fiber thickness thinned significantly on CD32 and CD55 relative to baseline, with no apparent sex differences (Fig. 2B–C).

**Figure 2:**
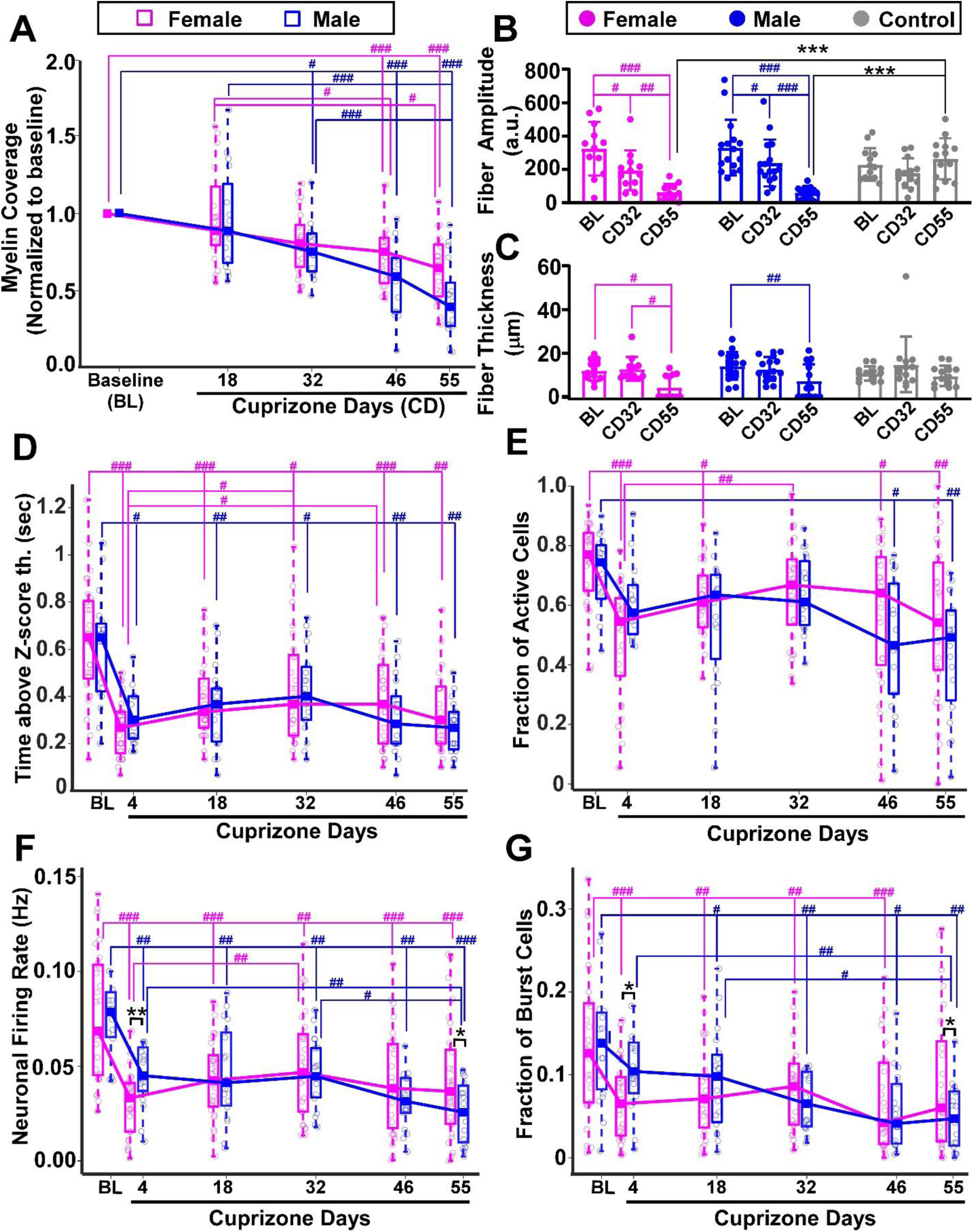
Sex-specific patterns of myelination coverage and CA1 activity suppression following Cuprizone consumption. **(A)** Myelin signal coverage, normalized to each mouse’s baseline levels, gradually declined during cuprizone consumption, becoming significantly lower than baseline after 32–46 days and showing more severe myelin loss in males than females on CD46–55 (n = 43 mice; 17 males, 26 females). Lines connect median values for each sex. For all Fig. 2 panels linear mixed-effects models with Tukey HSD corrections for post-hoc comparisons were used; #, P < 0.05; ##, P<0.01; ###, P < 0.001 for comparing different time points, and *, P < 0.05; **, P<0.01; ***, P < 0.001 for comparing across groups. Values of all statistical tests and p-values may be found in Tables S1 and S2. Boxplots show medians and 25–75% quartiles, and whiskers extend to the shorter of 1.5 × the interquartile range or the most extreme data point. Lines connect median values for each sex. **(B)** Same-fiber amplitude measurements showed a significant decrease between baseline, CD32, and CD55, with no sex differences (n = 19 mice, with 1-3 fibers measured per mouse; cuprizone group: 6 males, 6 females; control group: 3 males, 4 females). **(C)** Same-fiber thickness measurements showed a significant decrease between baseline and CD55, with no sex differences (same mice and fibers as in **B**). **(D)** Quantification of the average number of seconds per recording session, across all recorded neurons in each mouse, during which the fluorescence signal exceeded 3 z-scores revealed a significant decrease from baseline to CD4. This reduction remained significantly below baseline levels throughout the 55 days of cuprizone consumption in both male and female mice, with no sex differences. **(E)** Quantification of the fraction of neurons that fired at least one action potential in each recording, across all recorded neurons, showed a rapid decrease similar to that in panel D, followed by an intermediate recovery at CD32 and a second decrease at CD46–55. **(F)** CA1 mean firing rate showed significant decreases from baseline levels in both male and female mice. Females exhibited a significantly larger decrease than males on CD4, whereas males exhibited a significantly larger decrease than females on CD55. **(G)** The fraction of neurons exhibiting burst activity (at least 10 action potentials within a 1-s time window) showed a pattern similar to that in panel **F**, with a significantly larger decrease in females on CD4 and in males on CD55. Panels **D–G**: n = 48 mice (19 males, 29 females); females, 250–3525 neurons/mouse (median = 1083); males, 158–3207 neurons/mouse (median = 1178).

In contrast, neuronal activity changed much more rapidly following cuprizone exposure. Quantifying the average time per recording that neuronal fluorescence remained above a 3 z-score threshold (indicative of neuronal activity) revealed a steep, significant decrease of approximately 50% from baseline to the first recording on CD4 in both sexes, with no significant sex differences, and activity remained significantly below baseline throughout the cuprizone diet period (Fig. 2D). Extraction of AP firing patterns for all recorded neurons showed that the fraction of active neurons per recording also decreased significantly with cuprizone consumption, without significant sex differences (Fig. 2E).

Mean neuronal FR likewise decreased significantly from baseline to CD4 and remained significantly below baseline for the entire cuprizone period. Female mice showed significantly lower CA1 mean FR than males on CD4. After this initial decrease, CA1 FR partially recovered, reaching comparable levels in both sexes by CD18–32. A second decline then occurred during CD46–55 that was more pronounced in males, resulting in significantly lower mean FR in males than females by CD55 (Fig. 2F). Analysis of the fraction of neurons firing more than 10 APs within a 1-sec bin (burst cells) revealed a similar pattern: females showed a more severe decrease than males on CD4, whereas males showed a more severe decrease than females on CD55 (Fig. 2G). Normalizing data to each mouse’s own baseline showed a similar trend, with a significantly more severe decrease in the fraction of active neurons and mean FR in females on CD4, but no significant differences on CD55 (Fig. S2). Measuring the same activity parameters in the control group yielded no significant differences over time (Fig. S3).

### Acute cuprizone exposure has no apparent toxic effects on neurons

To test whether the short-term reduction in CA1 neuronal activity resulted from direct toxic effects of cuprizone on neurons, we performed whole-cell recordings from CA1 pyramidal neurons in acute hippocampal slices exposed *ex vivo* to different cuprizone concentrations. Following bath application of vehicle (0.3% dimethyl sulfoxide, DMSO), 50 µM cuprizone, or 100 µM cuprizone, we quantified the frequency–current (F-I) relationship, rheobase, resting membrane potential, and input resistance. None of these intrinsic properties differed significantly among the three groups (Fig. 3), indicating that acute exposure to cuprizone at concentrations up to 100 µM does not alter CA1 pyramidal neuron excitability under the conditions tested. These findings are consistent with previous work showing that cuprizone is selectively toxic to mature oligodendrocytes at concentrations as low as 25 µM^27^, and support the view that the early reduction in CA1 activity observed *in vivo* is driven by circuit- or glia-mediated mechanisms rather than by acute intrinsic toxicity to pyramidal neurons.

**Fig 3.**
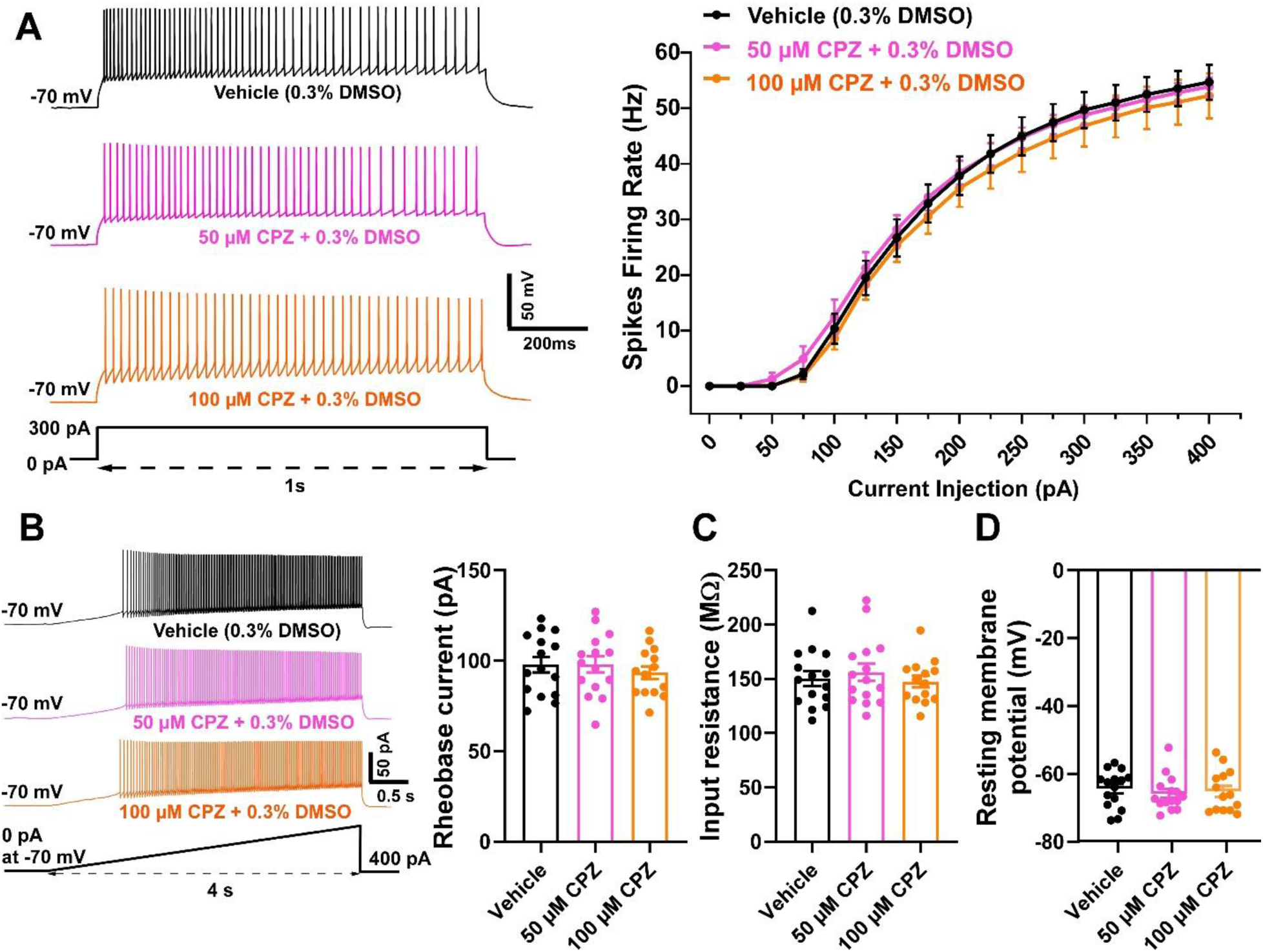
Acute cuprizone exposure does not alter CA1 pyramidal neuron excitability. **(A)** Left, representative action potential traces recorded from acute brain slices perfused with vehicle, 50 µM cuprizone, or 100 µM cuprizone. Right, frequency–current (F–I) curves showing similar firing rates in response to increasing current injections across the three groups (two-way mixed-design ANOVA with Bonferroni post hoc correction; F(2, 41) = 0.2739, P = 0.7618). Data are shown as the mean firing rate at each current step, and error bars indicate SEM. **(B)** Left, representative action potential traces recorded during the ramp current injection protocol used to determine rheobase current. Right, rheobase current was similar among the three groups (one-way ANOVA with Bonferroni post hoc test; F(2, 41) = 0.3904, P = 0.6793) **(C)** Input resistance was also similar among the three groups (one-way ANOVA with Bonferroni post hoc test; F(2, 41) = 0.4055, P = 0.6693). **(D)** Resting membrane potential did not differ among the three groups (one-way ANOVA with Bonferroni post hoc test; F(2, 41) = 0.2638, P = 0.7694). In panels **B–D**, bars represent the mean ± SEM, and each data point represents an individual recorded cell. The vehicle group included n = 15 cells from 7 mice (3 males, 4 females); the 50 µM cuprizone group included n = 15 cells from 4 mice (2 males, 2 females); and the 100 µM cuprizone group included n = 14 cells from 8 mice (4 males, 4 females).

### Remyelination treatment also restores CA1 activity

We next asked whether clemastine-enhanced remyelination could restore CA1 activity after prolonged demyelination. Following 55 days on the cuprizone diet, mice in the cuprizone group were returned to regular chow and randomly assigned to one of two treatment arms. The clemastine group received daily oral gavage of clemastine (10 mg/kg BW) for 45 days, and the vehicle group received oral gavage of vehicle solution alone. In agreement with previous studies, clemastine-treated mice showed enhanced remyelination compared with vehicle-treated mice^28,29^, as reflected by a monotonic increase in myelin coverage and significant increases in the signal amplitude and thickness of myelinated fibers (Fig. 4A-D). Notably, during the first 6-13 days after cuprizone cessation, myelin metrics increased modestly in both groups, reflecting spontaneous remyelination in the vehicle group. Thereafter, the vehicle group exhibited a significant monotonic decrease, whereas the clemastine group continued its increase over the remaining treatment period (Fig. 4B-D). At treatment day 45 (TD45), myelination in clemastine-treated mice had recovered to levels comparable to baseline, whereas vehicle-treated mice showed significantly lower myelinated-fiber signal amplitude than at their own baseline and than clemastine-treated mice at TD45 (Fig. 4E–Fx

**Figure 4:**
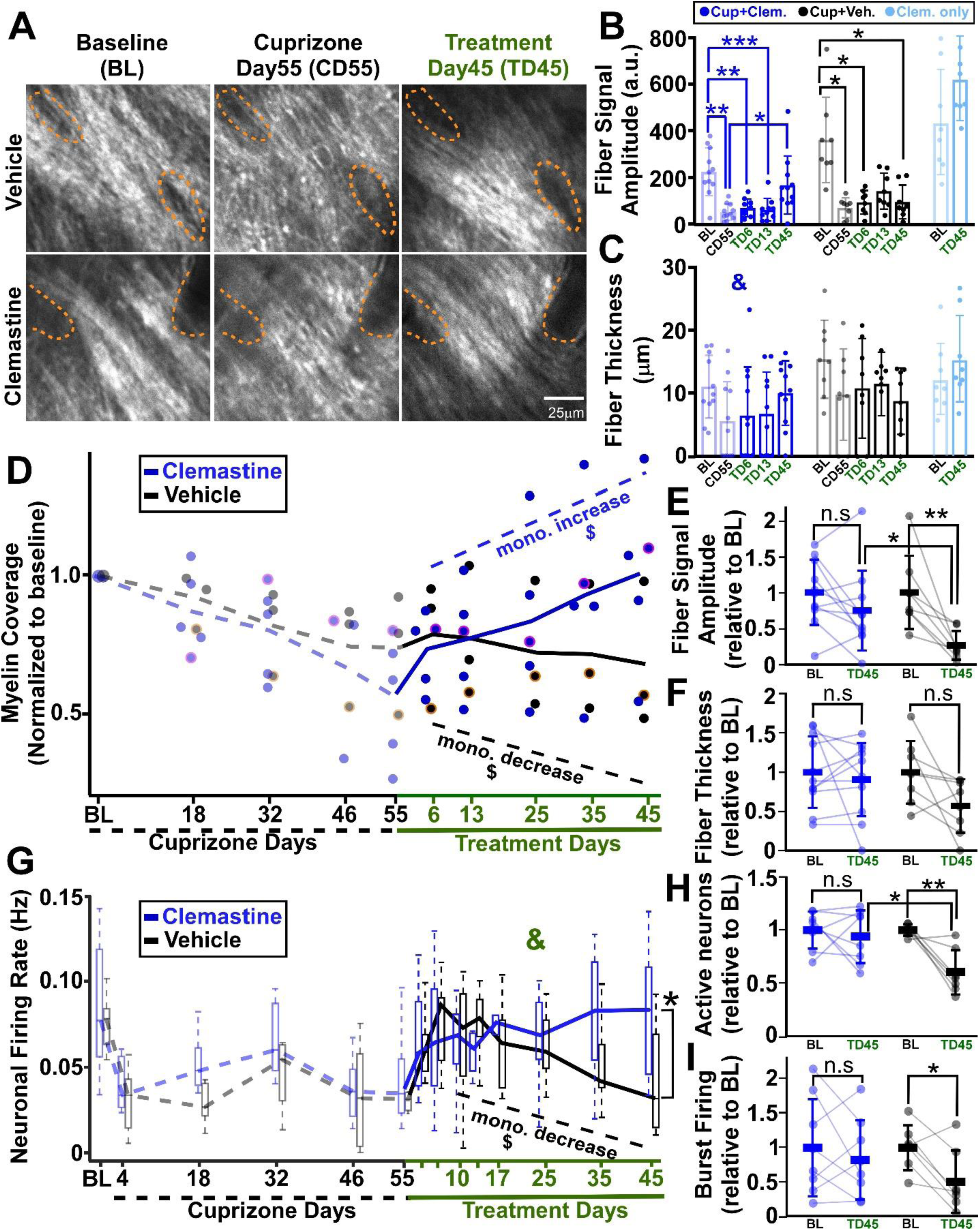
Clemastine treatment restores brain activity levels and enhances remyelination. **(A)** THG images of the same field of view at baseline, CD55, and TD45. Images are contrast-enhanced for visibility; orange dashed lines indicate blood vessels used to reliably re-identify the FOV. **(B)** Cross-sections of individual fibers were fitted with Gaussian functions to estimate changes in myelinated fiber signal amplitude over time (clemastine group: n = 11 fibers from 5 mice; vehicle group: n = 8 fibers from 3 mice; control group: n = 8 fibers from 5 mice). The control group was not fed cuprizone and received clemastine treatment. Both clemastine and vehicle groups showed significant decreases from baseline to CD55, whereas clemastine, but not vehicle, showed a significant increase during treatment. The vehicle group showed non-significant remyelination during the early treatment phase, followed by a late treatment phase decrease (Mixed-effects model with Tukey’s HSD correction for multiple comparisons; *, P < 0.05; \*\**, P* < 0.01*; **\*\*\****, P < 0.001 for comparisons across time points). **(C)** Estimation of the thickness of the same fibers shown in **B** revealed similar trends across vehicle and treatment groups, with a significant group effect (indicated by &, P<0.05) but no significant differences were found in the post hoc comparisons. For **B-C**, bars represent the mean ± std, and each data point represents thickness or amplitude of an individual fiber. Full statistical results and exact p values are provided in Tables S1 and S2. **(D)** Following treatment, mean myelin coverage monotonically increased in clemastine-treated mice and decreased in vehicle-treated mice (P = 0.028 for monotonic trends in both groups, Mann–Kendall test, indicated by $; n = 8 mice: 5 clemastine, 3 vehicle). Each dot represents data from one mouse; pale/dotted lines connect means during the cuprizone period and dark/solid lines connect means during the treatment period. Dots with magenta/orange borders correspond to the example data shown in **A**. **(E)** The myelinated-signal amplitude of the same fibers was compared between the baseline and TD45 recording sessions. In the clemastine group, signal amplitude did not differ significantly following treatment. In contrast, the vehicle group showed a significant reduction in signal amplitude at TD45 (P=0.127 and 0.007 for the clemastine and vehicle groups, respectively; same fibers as in B; two-tailed paired Student’s *t*-test). The clemastine group also showed a significant increase in signal amplitude at TD45 compared to the vehicle group (P=0.032, Student’s unpaired t-test). Bars indicate mean ± std. All values were normalized to the mean value from the corresponding baseline recording. **(F)** The thickness of the myelinated fibers, using the same fibers analyzed in **E**, was compared between the baseline and TD45 recording sessions. Consistent with the signal-amplitude measurements in **E**, fiber thickness did not decrease significantly in the clemastine group after treatment. The decrease in the vehicle group was not significant (P=0.605 and 0.059 for the clemastine and vehicle groups, respectively); same fibers as in **B** and **E**; two-tailed paired Student’s *t*-test). The difference between the clemastine and vehicle groups at TD45 was not significant (P=0.103, Student’s unpaired t-test). **(G)** Same-mouse CA1 FR was recorded longitudinally during cuprizone and treatment periods (pale/dotted and dark/solid lines/boxplots, respectively). During the first 6 treatment days, FRs increased in both groups. From TD13 to TD45, FRs decreased monotonically in vehicle-treated mice (P = 0.0275, Mann–Kendall test) but recovered toward baseline levels in clemastine-treated mice, resulting in a significant difference between the two groups at TD45 (P = 0.049; linear mixed-effects models with Tukey’s HSD, indicated by *; P = 0.027 for time × treatment interaction during the treatment period, indicated by &). n = 15 mice; 8 clemastine, 7 vehicle; clemastine: 321–3207 neurons per mouse, median 985; vehicle: 250–3021 neurons per mouse, median 915. Boxplots show medians and 25-75 quartiles, and the whisker length is the shorter of 1.5 times the inter-quartiles range or the extreme data point. **(H)** The proportion of neurons detected as active was compared between the baseline and TD45 recording sessions. Following treatment, the clemastine group showed no significant change, whereas the vehicle group showed a significant decrease (P=0.577 and 0.006 for the clemastine and vehicle groups, respectively, two-tailed paired Student’s *t*-test, same mice as in **G**). The clemastine group also showed a significant increase at TD45 compared to vehicle group (P=0.015, Student’s unpaired t-test). All values were normalized to the mean value from the corresponding baseline recording. **(I)** Consistent with H, the proportion of burst-firing neurons was compared between the baseline and TD45 recording sessions. The clemastine-treated group showed no significant change at TD45 relative to baseline, whereas the vehicle-treated group showed a significant decrease (P=0.475 and 0.023 for the clemastine and vehicle groups, respectively, two-tailed paired Student’s *t*-test, same mice as in **G**). The difference between both groups at TD45 was not significant (P=0.26, Student’s unpaired t-test). All values were normalized to the mean value from the corresponding baseline recording. Full statistical results and exact P values for panels **B-D,G** are provided in Tables S1 and S2.

CA1 FR followed a similar time course, but with larger amplitude changes. Both groups showed a transient increase during the first 6 days of treatment, followed by a significant monotonic decline in the vehicle group and a gradual increase in the clemastine group. These divergent trajectories resulted in a significant effect of clemastine treatment on the fraction of CA1 neurons that were active and their average FR at TD45 (Fig. 4G-H). In addition, at TD45, activity metrics in the clemastine group recovered to levels comparable to same-mouse baseline values, whereas the vehicle group showed significant decreases in most metrics relative to baseline (Fig. 4H–I; Fig. S2E–F).

Because fewer mice completed both the cuprizone and treatment phases than completed the cuprizone diet period alone, we tested whether this subset showed survival bias in body weight, myelination status, or CA1 activity levels. No relevant differences were found between these groups at baseline and CD55 (Fig. S4). In addition, in control mice that did not receive cuprizone, clemastine treatment produced a non-significant trend toward increased myelin coverage and fiber signal amplitude and thickness, but no trends or significant changes in CA1 activity patterns (Fig. 4B–C, Fig. S3). These stable activity patterns were similar to CA1 activity patterns in mice that were implanted with hippocampal windows and received regular diet without additional intervention (Fig. S3). Together, these data indicate that clemastine treatment enhances hippocampal remyelination and restores CA1 activity to pre-cuprizone levels (Fig. 4E). This protective effect appears to preferentially target circuits disrupted by prior demyelination.

### Cellular-level responses to treatment

In a subset of the cuprizone group, we longitudinally tracked the same neurons across the entire imaging period (n = 6 mice; 3 treated with clemastine and 3 with vehicle), and compared same-cell activity across baseline, CD55, TD10, and TD45. No spatial organization of neurons with increased or decreased firing was identified within the recorded fields of view (Fig. 5A). We next examined whether cellular properties, such as baseline FR, were related to the early and late recovery patterns defined in Fig. 4E (early phase: CD55-TD10; late phase: TD10-TD45). During the early treatment phase, changes in neuronal FR in the clemastine group showed only a weak dependence on baseline FR, with neurons with lower baseline FR exhibiting slightly greater recovery of their mean FR. By contrast, in vehicle-treated mice, baseline FR was positively correlated with the increase in FR, such that neurons with higher baseline FR showed larger increases in mean FR between CD55 and TD10 (Fig. 5B, Fig. S5). During the late treatment phase, the relationship between baseline FR and treatment response reversed between the two groups: in clemastine-treated mice, neurons with higher baseline FR showed larger increases in FR, whereas in vehicle-treated mice, neurons with higher baseline FR showed larger decreases (Fig. 5C–E), and the variability of these changes also increased with baseline FR (Fig. 5F). Together, these findings suggest that cuprizone withdrawal leads to a transient increase in neuronal activity in vehicle-treated mice, with neurons of higher baseline activity increasing more, followed by a decline phase in which those same neurons decrease more strongly, whereas clemastine treatment stabilizes neuronal activity during the early treatment phase and enhances it during the late treatment phase, opposing the trajectory observed in the vehicle group.

**Figure 5:**
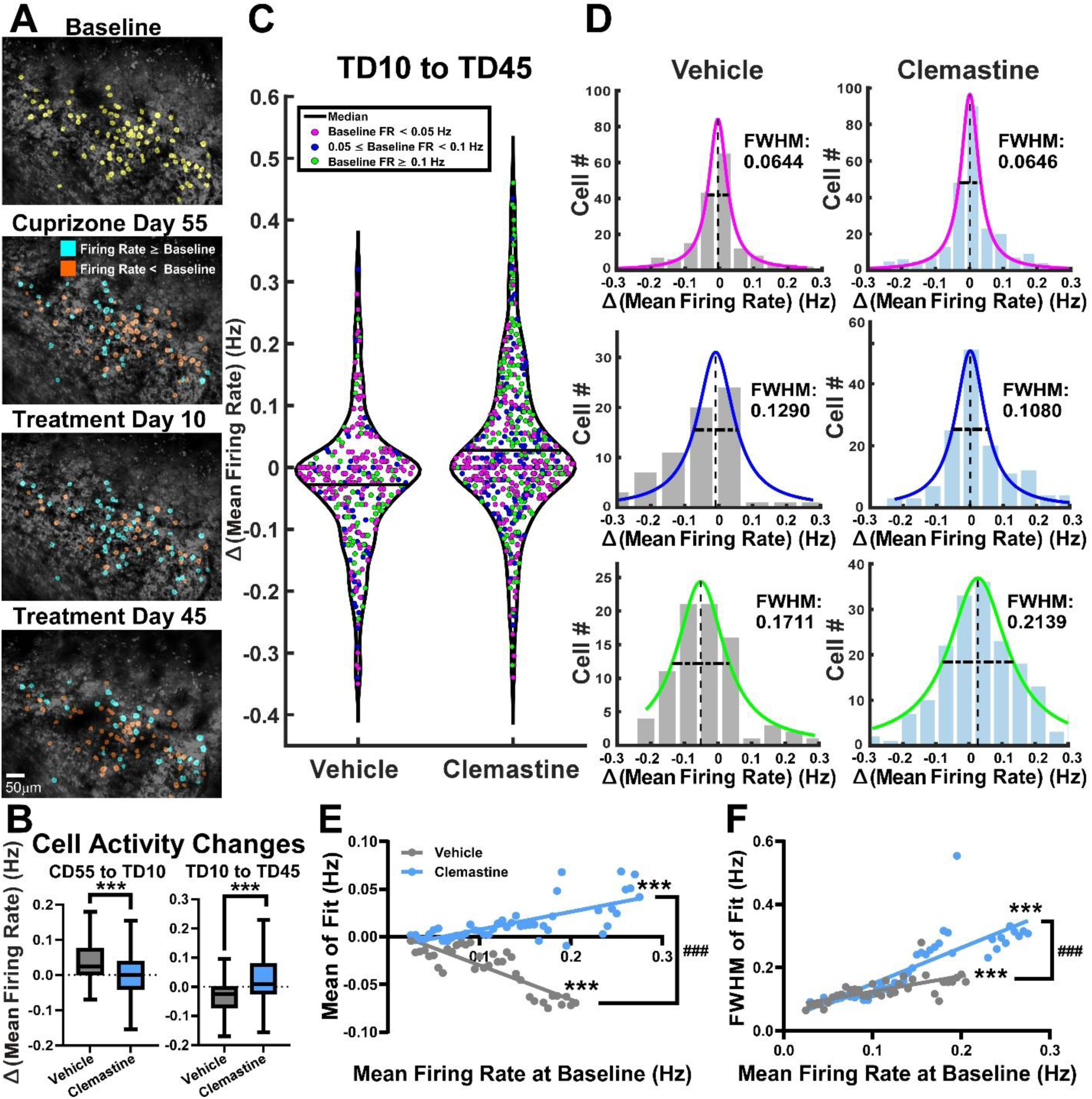
Single-cell correlates of recovery during clemastine treatment. **(A)** Example tracking of the same CA1 neurons at baseline, CD55, TD10, and TD45 (n = 87 cells, highlighted in yellow in the baseline image). Cyan- and orange-labeled cells exhibit higher and lower mean firing rates than their baseline levels, respectively. **(B)** Distributions of changes in same-cell mean FRs from CD55 to TD10 (early treatment phase, left) and from TD10 to TD45 (late treatment phase, right) for vehicle and clemastine groups (n = 6 mice; 3 per group; 165 and 336 tracked cells in vehicle and clemastine groups, respectively). The vehicle group showed a larger increase in FR than the clemastine group during the early treatment phase, whereas the clemastine group showed a larger increase than the vehicle group during the late treatment phase (P < 0.0001 for both comparisons, unpaired two-sided Student’s t-test). **(C)** Violin plot of changes in same-cell firing rate during the late treatment phase, with dots colored according to low, medium, or high FRs at baseline (vehicle: 306 cells; clemastine: 553 cells from the same 3 mice per group as in **B**) **(D)** Histograms of low-, medium-, and high-baseline firing-rate cells from **C** fitted with Lorentzian functions. Neurons with high baseline activity tended to have larger FWHM values, with positive mean values in clemastine-treated mice and negative mean values in vehicle-treated mice. **(E)** Cells from **C** were grouped into partially overlapping baseline firing-rate bins, and their histograms were fitted with Lorentzian functions as in **D**. Lorentzian means were positively correlated with baseline firing rate in clemastine-treated mice and negatively correlated in vehicle-treated mice (clemastine: Pearson’s r = 0.7024; vehicle: r = −0.8451; ***, P < 0.001 for both regression lines; ###, P < 0.001, ANCOVA for detecting differences in regression line slopes). **(F)** In both groups, variability (FWHM) increased with baseline FR (clemastine: Pearson’s r = 0.8436; vehicle: r = 0.6602). Additional details are provided in the Supplementary Statistical Data.

## Discussion

In this study, we used an integrateddual-imaging approach to simultaneously track neuronal activity and myelination status throughout cuprizone-induced demyelination and clemastine-mediated remyelination in mice by combining TPLSM recording of GECI signals with label-free THG imaging of myelin. We report three major findings. First, neuronal firing suppression occurs as early as 4 days after cuprizone initiation and precedes myelin loss by weeks, revealing a critical temporal dissociation between functional and structural pathology. Second, neuronal firing rates show a biphasic decline over the cuprizone consumption period, with early activity loss that is more severe in females and late-stage decline that is more severe in males, revealing previously unreported sex-specific functional vulnerabilities. Third, clemastine treatment not only promoted remyelination but also restored neuronal activity toward baseline levels, with the magnitude and direction of single-neuron responses associated with baseline firing rate.

Previous work from our group demonstrated that cuprizone consumption significantly reduces CA1 activity within 9 days, well before peak demyelination at 3–5 weeks^8,13^. The present study extends these findings by examining earlier time points, linking longitudinal brain-activity recordings with myelin measurements in the same mice, and using a substantially larger sample to evaluate potential sex differences. We detected reduced CA1 activity as early as 4 days after cuprizone initiation (Fig. 2D–G), substantially preceding the first detectable myelin loss at CD32 (Fig. 2A–C). Importantly, acute cuprizone exposure did not alter neuronal intrinsic excitability, even at concentrations up to 100 μM (Fig. 3). Although cuprizone has been reported to reduce oxygen consumption in cultured cortical neurons after 24 h of exposure, glial cells were more susceptible to cuprizone-induced mitochondrial dysfunction and loss of viability in that study^30^. Our findings argue against an acute, direct effect on neuronal intrinsic excitability as the primary explanation for the early reduction in CA1 activity. Moreover, a prior electrophysiological study in cuprizone-fed mice found no acute change in CA1 synaptic transmission, but instead observed a gradual decline that became significant after 3–4 weeks^13^. Consistent with a glial contribution to this early phenotype, mature oligodendrocytes are selectively vulnerable to cuprizone, exhibiting mitochondrial dysfunction and reduced viability after 24 h of exposure at concentrations as low as 25 μM^27^. Together, these findings indicate that acute, direct cuprizone effects on neuronal intrinsic excitability are unlikely to be the principal driver of the early CA1 activity loss. Instead, the rapid reduction in CA1 activity is more consistent with early oligodendrocyte dysfunction and its downstream circuit consequences, although direct cuprizone-induced metabolic effects on neurons may also contribute.

Interestingly, previous studies identified hyperactivity in mice both during cuprizone administration and after its cessation. For example, Bacmeister et al. showed increased activity in motor cortex neurons shortly after cessation of a 3-week cuprizone diet, which returned to control levels over the following two weeks^31^. This time course aligns with the transient increase in activity we identified during the early treatment phase (Fig. 4E), which was present in both the vehicle and clemastine groups. Notably, the mean increase during this early phase was higher in the vehicle-treated group, and its cellular impact differed by baseline firing rate: clemastine-treated mice showed slightly stronger increases in low-baseline-FR neurons, whereas vehicle-treated mice showed stronger increases overall than clemastine-treated mice, driven specifically by high-baseline-FR neurons (Fig. S5C). This finding suggests that clemastine may have a protective effect against the early exaggerated hyperactivity of high-baseline-FR neurons following cessation of cuprizone consumption. In another study, Hamada and Kole showed that layer V neurons undergo changes in cellular and axon initial segment properties after 5 and 9 weeks of cuprizone diet, reflected in increased burst activity and cellular excitability^32^, which were not identified in the current study. Several factors may contribute to these differences. First, Hamada and Kole’s findings were acquired *ex vivo* using whole-cell recordings from layer V neurons and may therefore reflect differences in circuit-level impact and recording modality relative to our *in vivo* TPLSM recordings of CA1 neurons using GECIs. Second, baseline myelination levels and the extent of cuprizone-induced myelin loss differ between pyramidal CA1 and layer V neurons^26^. In addition, cuprizone has been shown to affect PV⁺ interneurons and circuit-level activity patterns^33^, an effect that may not be fully captured in slice electrophysiology studies. Finally, Hamada and Kole’s work did not examine early time points, such as CD4, that we identified in our study.

The temporal dissociation between functional and structural changes indicates that neuronal dysfunction in the cuprizone model is not simply a consequence of demyelination but likely reflects earlier disruptions in neuron-oligodendrocyte metabolic coupling, synaptic transmission, or inflammatory signaling. Several mechanisms may account for this early activity loss. First, cuprizone primarily targets mature oligodendrocytes through copper chelation and mitochondrial dysfunction, causing cellular stress and early death before overt demyelination^27,34,35^. Stressed oligodendrocytes may fail to provide metabolic support to axons and thus disrupt neuronal energy homeostasis. Second, oligodendrocyte dysfunction and apoptosis may trigger microglial activation and inflammatory cytokine release^34,36,37^ within the first week of cuprizone exposure, which could directly suppress synaptic transmission^38^. Third, subtle changes in myelin structure or composition are likely to occur before gross demyelination becomes detectable by THG microscopy, altering axonal conduction velocity and excitability through changes in ion channel distribution or axon initial segment properties^10,39–41^.

Sex differences in the cuprizone model have been reported in several animal species^42–44^, including C57BL/6J^45,46^, which underlies the Thy1-jRGECO1a line used in this study. Our findings on sex differences in myelin coverage, but not in individual fiber characteristics (Fig. 2A-C), are in line with prior reports of similar demyelination patterns in male and female C57BL/6 mice^45,46^. However, a new finding we report is that the timing and severity of neuronal dysfunction differ markedly between sexes. Female mice showed significantly lower CA1 firing rates than males at day 4 of cuprizone treatment (Fig. 2F-G), suggesting greater early vulnerability to oligodendrocyte stress. In contrast, males exhibited more severe activity decline during the late cuprizone phase (CD46-55), which paralleled the greater myelin coverage loss in males (Fig. 2A). These biphasic, sex-divergent patterns suggest that males and females engage different compensatory mechanisms or have distinct thresholds for neuronal activity impairment during demyelination. The early sex difference is particularly notable because it occurs well before detectable myelin loss and remains evident when data are normalized to same-mouse baseline levels (Fig. S2A-D; note that normalization to each animal’s own baseline accounts for differences in baseline measurements across animals but may amplify noise when baseline values are small). This temporal pattern of sex-specific vulnerability may have important translational implications for MS and other demyelinating diseases. MS shows a 3:1 female predominance in incidence, yet paradoxically, men often develop more severe progressive disease with greater neurodegeneration and faster disability accumulation^47,48^. Our finding that females exhibit earlier functional impairment, which precedes the earliest signs of myelin loss, mirrors aspects of this clinical pattern, suggesting that sex differences in MS progression may reflect intrinsic differences in how neuronal circuits respond to oligodendrocyte injury or stress. Multiple mechanisms likely contribute to these divergent patterns, including hormonal modulation of neuronal excitability and metabolic stress responses, sex chromosome effects on oligodendrocyte function, and sex-dependent differences in microglial activation and inflammatory cytokine production^43,49–52^. Understanding the cellular and molecular basis of these sex-specific vulnerabilities could motivate the development of remyelination therapies that are better tailored to male and female patients.

Remyelination therapies hold promise for slowing or reversing disability in MS, particularly during progressive stages of disease^53^. Clemastine, an FDA-approved antihistamine, promotes oligodendrocyte maturation and myelin repair and has shown efficacy in both preclinical models and the ReBUILD clinical trial^20,28,29^. Our findings demonstrate that clemastine not only accelerated remyelination (Fig. 4B–F) but also restored CA1 neuronal firing rates close to baseline levels, whereas vehicle-treated mice showed continued decline (Fig. 4G-I, Fig. S2E-F). This functional restoration extended beyond simple recovery of population-level activity: at the single-neuron level, clemastine preferentially enhanced firing in neurons with higher baseline activity, whereas in vehicle-treated mice, high-baseline neurons showed a greater decline (Fig. 5E). This baseline-activity-associated effect suggests that neuronal responses to clemastine differ according to pre-existing functional properties, potentially including excitability, synaptic connectivity, metabolic demand, or myelination status. Neurons with higher baseline firing rates are more metabolically demanding and thus may be more vulnerable to loss of oligodendrocyte metabolic support, making them particularly sensitive to their remyelination status. Alternatively, highly active neurons may provide stronger pro-myelination signals through activity-dependent release of neurotrophic factors^54^, creating a positive feedback loop with clemastine-enhanced oligodendrogenesis. These neurons may also be more myelinated than low-activity neurons before cuprizone administration, and clemastine may restore them to their pre-cuprizone conditions. These findings align with growing evidence that neuronal activity and myelination are bidirectionally coupled^31,54,55^ and suggest that remyelination therapies may confer dual benefits, promoting recovery of mature oligodendrocytes and structural repair of myelin, as well as functional restoration of neuronal circuits. In light of recent clinical trial data indicating potential adverse effects of clemastine in progressive MS^56^, it is notable that clemastine treatment in control mice not exposed to cuprizone did not alter neuronal firing rates (Fig. S3). The absence of detectable firing-rate changes in clemastine-treated control mice suggests that the effects observed after cuprizone exposure do not reflect a generalized effect of clemastine on CA1 neuronal activity but depend on the pathophysiological context produced by cuprizone exposure.

This study’s longitudinal dual-imaging approach enabled us to track structure-function relationships in the same animals over time, reducing inter-animal variability and increasing statistical power^57–60^. To our knowledge, this is the first study to simultaneously monitor neuronal activity and myelination at cellular resolution throughout demyelination and remyelination *in vivo*. This approach could be extended to other demyelination models or genetic models of neurodegeneration and could facilitate preclinical testing of novel remyelination therapies by providing longitudinal functional readouts alongside structural measures. Several limitations should be acknowledged when interpreting our findings. First, the complexity of longitudinal imaging limited our sample size, preventing us from fully characterizing sex differences during the remyelination phase. Second, we examined only a single clemastine dose, based on previous reports^28,29^; dose-response studies are needed to identify optimal dosing regimens. Third, while we focused on CA1 pyramidal neurons, future work should examine other brain regions and cell types to determine whether the rapid activity loss and sex-specific vulnerabilities we observed are generalizable. Finally, our study design prioritized population-wide changes in firing patterns and myelination and therefore could not directly link the activity of a specific neuron to the myelination level of its axon. Future work combining cellular-resolution activity measurements with axon-specific myelin monitoring will be essential to resolve this relationship.

## Materials and methods

All surgical and experimental procedures were conducted in compliance with protocols approved by the Institutional Animal Care and Use Committee and Institutional Biosafety Committee of the Cleveland Clinic Research. The ARRIVE guidelines were used to report the below findings^61^.

### Surgical procedures

Thy1-jRGECO1a mice (10-12 weeks old) were implanted with hippocampal windows above their left hippocampus following published protocols^62,63^. Briefly, mice were anesthetized using isoflurane (3% for induction, 1.5% during surgery), the scalp skin was removed, and a circular craniotomy was drilled through the skull (3.2mm diameter, −2/-2 mm AP/ML from Bregma, OMNIDRILL35, WPI). The cortex and corpus callosum tissue above the hippocampus were aspirated, while leaving the hippocampus intact. A glass coverslip (3mm diameter #1 glass, Warner Instruments) attached to a metal cannula (2.65/3.2mm internal/outer diameters, 1.8mm length, New England Small Tube Company, 304L stainless steel) was placed above the hippocampus. The cannula and a headbar were secured to the skull using dental cement (Contemporary Ortho-Jet) and a cyanoacrylate glue.

### Longitudinal recordings

Mice were intramuscularly injected with Chloroprothixene Hydrochloride (30µl of 0.33mg/ml, Spectrum Chemical) and were kept on a 37°C heating pad under light anesthesia (0.5-0.75% isoflurane)^13^. TPLSM was used to record spontaneous activity of CA1 pyramidal neurons expressing jRGECO1a using 1100 nm excitation light (100-200mW, Insight X3, Spectra-Physics), a 10x 0.5 NA objective (TL10X-2P, Thorlabs), and a Bergamo II microscope (Thorlabs) equipped with resonant/galvo scanners and a GaAsP PMT detector (512×512 pixels, 600×600μm^2^, 30 frames/sec, 607/70nm emission filter, Semrock). Three FOVs were typically collected for 200 seconds each. One to two baseline recordings were conducted before the initiation of cuprizone diet, as well as on cuprizone days (CDs) 4, 18, 32, 46, and 55, and subsequently on treatment days (TDs) 3, 6, 10, 13, 17, 25, 35, and 45. Recording days were selected to capture the expected early changes at the beginning of cuprizone and treatment periods, and then to monitor the longitudinal changes. All recording sessions were performed identically within the mouse CA1 region. For a subset of mice (n=6) the same neurons were recorded across sessions and we verified their identity using the CellReg V1.5.5 algorithm^64^. For FOVs that passed the CellReg test, we included cells with P_same_>0.5 values.

THG recordings were conducted using the same microscope using low-repetition-rate laser at 1260 nm excitation wavelength (15-60 mW, Carbide 40W laser and Orpheus-F OPA, Light Conversion, 1MHz repletion rate) using galvo-galvo scanners and a GaAsP PMT detector (1024×1024 pixels, both 200×200 and 400×400 μm^2^ FOVs were acquired, ∼1-2 frames/sec, 434/32nm detection filter, Semrock). To minimize the risk of heat-induced damage, recordings were conducted once during baseline, and on CDs 18, 32, 46, and 55, and then on TDs 6, 13, 25,

35, and 45.

### Mouse model of demyelination and remyelination

Mice were kept in standard housing (14-hour light/10-hour dark cycle, *ad libitum* access to food and water) and were allowed 4–7 weeks of recovery after craniotomy before recordings began. After initiating recordings in a subset of mice, we found that a minimum of 40 days was required to achieve full recovery, as reflected by stable CA1 activity. Accordingly, only mice with at least one baseline recording obtained 40 or more days post-craniotomy were included in the study; the remaining mice were excluded (n = 25, Fig. S6). Following baseline recordings, mice were randomly assigned to one of two groups: the cuprizone group, fed a 0.3% cuprizone diet (TD.140805, Envigo), or the control group, maintained on a regular diet either alone or in combination with clemastine treatment (see below). Cuprizone pellets were stored at 4°C and replaced three times per week. After 55 days on the cuprizone diet, mice were returned to a regular diet for an additional 45 days while receiving either clemastine or vehicle treatment until the study endpoint.

Mice that completed the cuprizone exposure period, as well as control mice assigned to receive clemastine, were randomly allocated to daily treatment with either clemastine or vehicle (Sigma-Aldrich; 10 mg/kg body weight). Mice were orally gavaged once daily with either a solution of 1.33 mg/ml clemastine fumarate in 90% H₂O, 5% Tween-20, and 5% DMSO, or with vehicle solution alone. Mice removed from the study during the cuprizone or treatment phases owing to health concerns or poor cranial window quality were excluded from the analysis (n = 36, Table S3). No blinding was used.

### Brain slice preparation and *ex vivo* brain slice recordings

C57BL/6J mice aged 4–5 weeks were anesthetized with 2.5% isoflurane, after which the brains were rapidly removed and sectioned in an ice-cold choline chloride-based cutting solution containing (in mM): 110 choline chloride, 24 NaHCO₃, 20 D-glucose, 1.2 NaH₂PO₄, 2.5 KCl, 7 MgCl₂, 5 HEPES, 0.5 CaCl₂, 2 thiourea, 3 sodium pyruvate, and 5 sodium ascorbate. Coronal brain slices (300 μm thick) were prepared using a vibratome (Leica VT1200). The slices were transferred to artificial cerebrospinal fluid (ACSF) maintained at 34°C for 30 min. The ACSF contained (in mM): 124 NaCl, 24 NaHCO₃, 12.5 D-glucose, 1.2 NaH₂PO₄, 2.5 KCl, 2 MgCl₂, 5 HEPES, and 2 CaCl₂. All solutions were continuously bubbled with 95% O₂ and 5% CO₂, maintained at pH 7.3–7.4, and adjusted to an osmolality of 300–310 mOsm/kg. The slices were subsequently transferred to ACSF at room temperature until electrophysiological recordings.

DMSO was used as the vehicle to dissolve cuprizone (CPZ; bis(cyclohexanone)oxaldihydrazone, 14690; Sigma-Aldrich). The control group received ACSF containing 0.3% (v/v) DMSO, and the same DMSO concentration was maintained in the 50 μM and 100 μM CPZ treatment groups. Before recording, slices from all experimental groups were continuously perfused with their respective solutions for at least 20 min. CA1 pyramidal neurons in the dorsal hippocampus were visually identified using an upright microscope (BX51WI, Olympus) equipped with a 40× water-immersion objective. During recordings, slices were continuously perfused with ACSF at a flow rate of 2–4 mL/min and maintained at 32–34°C. Borosilicate glass patch pipettes (1.5 mm outer diameter, 0.86 mm inner diameter) with a tip resistance of 3–4 MΩ were pulled using a P-97 micropipette puller (Sutter Instruments) and filled with a K-gluconate-based internal solution containing (in mM): 150 K-gluconate, 4 KCl, 10 HEPES, 2 MgCl₂, 0.5 EGTA, 4 Mg-ATP, 0.3 Na-GTP, and 10 phosphocreatine di(tris) salt. The internal solution was adjusted to pH 7.2–7.3 and an osmolality of approximately 290 mOsm/kg.

Whole-cell patch-clamp recordings were obtained using a Multiclamp 700B amplifier (Molecular Devices) and digitized with a Digidata 1550A (Molecular Devices). Signals were sampled at 20 kHz and low-pass filtered at 10 kHz. Action potential firing was evoked by injecting depolarizing current steps in 25 pA increments up to 400 pA. Rheobase was determined by holding neurons at −70 mV and applying a 4 s current ramp from 0 to 400 pA. The current at which the first action potential was elicited was defined as the rheobase. Recordings in which the series resistance changed by more than 20% during the experiment were excluded from analysis.

### Data analysis

Analyses were based upon custom MATLAB scripts (MathWorks), similar to our previous work^13^. Raw fluorescence movies were registered in order to correct for small movements^65^. Somatic regions of interest (ROIs) were segmented using CellPose^66^. Pixels of each ROI were averaged to calculate the time-dependent fluorescence signal (F). Baseline fluorescence (F_0_) was estimated as the median of F and z-score traces were calculated as (F-mean(F)/std(F)). AP numbers and timing were estimated using the SpikeML algorithm using jRGECO1a published parameters^15,22^. Myelin coverage was quantified by classifying the raw images to 3 levels (myelin signal, bright background, and dim background) using Otsu’s method to find the threshold values and calculating the ratio of the brightest pixels (myelin signal) vs. all pixels. Myelin fiber properties were measured by fitting a Gaussian function to cross-sections of the same visually-identified fibers across different recording dates. Thickness was defined as the fit’s full width at half maximum (FWHM). The amplitude was defined as the maximum of the fit minus its minimum. In case the Gaussian fit to the fiber cross-section yielded a flat line, such as in fibers that were barely visible on CD55, the thickness was set to 0 and the amplitude was calculated without modifications. All recordings were inspected for quality issues, like movement artifacts and relocation of the same myelinated fibers. In case issues were found, the specific session was excluded (TPLSM recordings: n_1_= 9 sessions from n_2_= 9 mice; THG recordings: n_3_= 22 sessions from n_4_= 18 mice, Table S4).

Data from the same-cell analysis were compared across four recording days: baseline, CD55, TD10, and TD45. The same-cell FRs were calculated, and outliers were excluded using the ROUT method (Q = 0.01). Separately, cells which were identified just in the [baseline, TD10, and TD45] or [baseline, CD55 and TD10] datasets, were also excluded of outliers (ROUT, Q = 0.01) and binned into partially-overlapping bins of ±0.025 Hz baseline FRs. We calculated histograms of change in average FR from TD10 to TD45 or CD55 to TD10 for each bin, which were fitted with scaled Lorentzian functions: *Y* = *a*/(1 + ((*x* − *b*)/*γ*)^2^). Bins with less than 20 cells or R^2^ of fit lower than 0.8 were omitted. Linear regressions of the baseline FRs vs. the function’s peak and FWHM were calculated to identify correlations between the parameters. Electrophysiological data were analyzed using Clampfit 11.2 (Molecular Devices). Representative traces were prepared using Igor Pro 8.04 (WaveMetrics).

### Statistical analyses

Analyses of TPLSM and THG data were conducted using GraphPad Prism (GraphPad Software, version 11.0.2). Average FRs and myelin signals were fit with mixed-effects models. The mixed-effects models used a compound symmetry covariance matrix and were fit using Restricted Maximum Likelihood. In the absence of missing values, this method gives the same P values and multiple comparisons tests as repeated-measures ANOVA. Data fit was confirmed by checking the respective QQ plots. We pursued *post hoc* comparisons only if the respective mixed-effects models’ P values were significant (i.e. *post hoc* comparisons between different time points were considered only if the mixed-effects models’ p value for time effect was significant. Significant P value for time x group interactions led to *post hoc* comparisons of both). Tukey’s HSD correction was applied for multiple comparisons. For baseline-normalized analyses, we excluded the baseline data and used the same mixed-effects models with Tukey’s HSD for the remaining of the data. Significance with respect to the baseline data was tested using a one-sample Student’s t-test, followed by Bonferroni corrections for multiple comparisons. Mann-Kendall test was used to detect monotonic trends between mean data points across different recording dates.

Statistical analyses of the electrophysiological data were performed using GraphPad Prism 11.0.2). Data were modeled using one-way and two-way ANOVA with Bonferroni correction for multiple comparisons.

A significance threshold of P < 0.05 was used for all statistical tests. Details on statistical effect sizes may be found in Supplementary Tables S1-S2.

## Data availability

Data related to this study will be available from the corresponding author upon reasonable request.

## Acknowledgements

The authors would like to thank Drs. Christopher Nelson and Xuefeng (Chris) Liu for helpful comments and suggestions, as well as to Drs. Monica Boyle, Kevin Dines, and Mrs. Iliana Ruiz for input on the study design. This study was supported by Bristol-Myers Squibb. The funder had no role in the data collection or analysis, decision to publish, or preparation of the manuscript.

## Author contributions

H.D. and B.T. conceived the project. H.D. and A.D. designed the study and interpreted the results. A.D. performed all surgeries. A.D., J.B. and J.I. performed optical recording experiments. A.D., H.T., J.B., S.B., J.I., G.T., P.A., S.S., J.M., P.I. and W.L. contributed to the optical data analysis. H.D., T.M.D. and R.K. designed the electrophysiological recordings experiments, R.K. performed the experiments and the respective data analysis. A.D., and H.D. wrote the manuscript with comments from all authors.

## Supplemental information

Document S1. Supplemental Figures S1-S6 and Tables S1-S4

