## Supplementary Figures and Tables for "Remyelination therapy restores suppressed neuronal activity in a mouse model of demyelination"

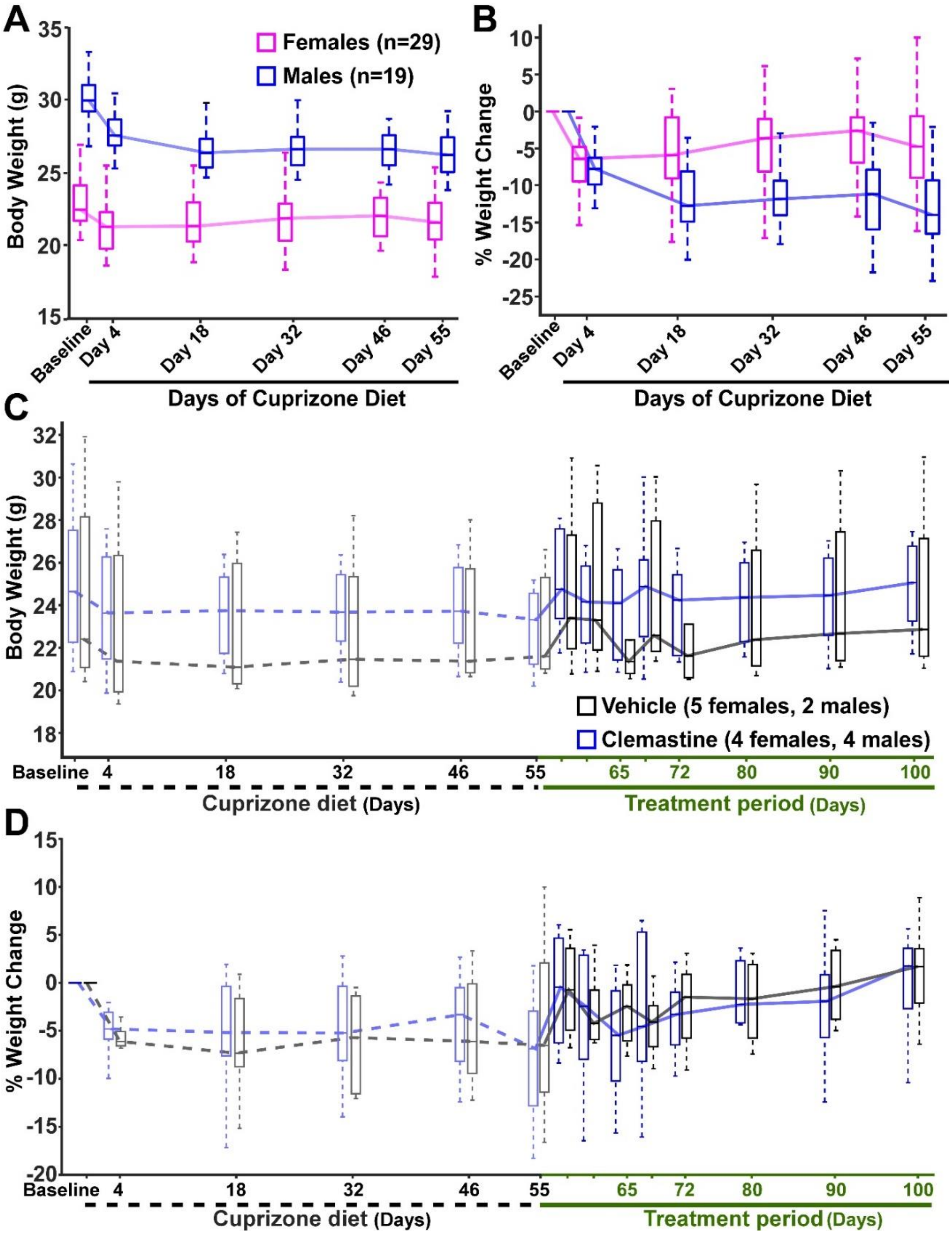


**Figure S1, Changes in body weight during the cuprizone diet and treatment periods, related to Figure 1**

**(A)** Both male and female mice lost body weight during the cuprizone diet period (n = 48 mice; 19 males, 29 females). Time effect: $F(2.758, 126.3) = 51.65$, $P < 0.0001$; group effect: $F(1, 46) = 78.63$, $P < 0.0001$;

time × group interaction: $F(5, 229) = 15.09$, $P < 0.0001$ (mixed-effects model with restricted maximum likelihood and Tukey's HSD *post hoc* correction for multiple comparisons).

**(B)** Male mice lost a larger percentage of their initial body weight than females (at CD55, median body weight loss from baseline was 13.99% and 4.74% for males and females, respectively; same mice as in **A**).

**(C)** During the treatment period, both the Clemastine and vehicle groups showed a gradual increase in body weight.

**(D)** Same data as in **C**, normalized to each mouse's initial body weight. Both the Clemastine and vehicle groups showed a similar increase in body weight.

Boxplots show the $25^{th}$-$75^{th}$ percentile range, midlines correspond to the median of each distribution and are connected with solid lines, and the whisker length is the shorter of 1.5 times the 25th–75th range or the extreme data point. Outliers are not shown.

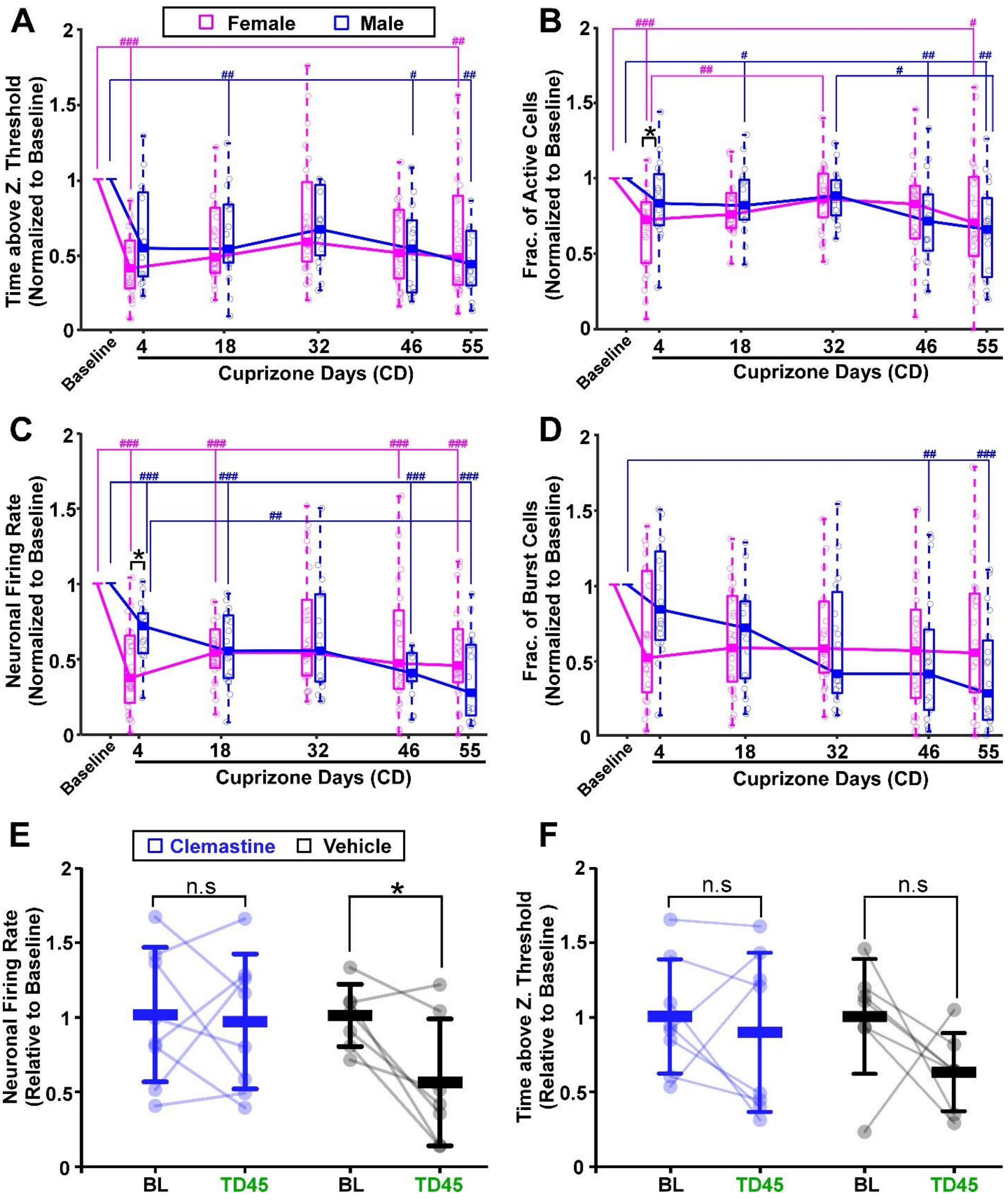
A
Female
Male
Time above Z. Threshold (Normalized to Baseline)
Baseline
Cuprizone Days (CD)
B
Frac. of Active Cells (Normalized to Baseline)
C
Neuronal Firing Rate (Normalized to Baseline)
D
Frac. of Burst Cells (Normalized to Baseline)
E
Clemastine
Vehicle
Neuronal Firing Rate (Relative to Baseline)
n.s
BL
TD45
F
Time above Z. Threshold (Relative to Baseline )

**Figure S2, Normalized data show sex-specific patterns of CA1 activity suppression following Cuprizone consumption, related to Figures 2 and 4**

(**A–D)** Panels A-D show the same data as in Figure 2D–G, with all values normalized to same-mouse mean baseline values. Similar to Fig. 2F, panel **C** shows a larger decrease in mean firing rate at CD4 in female mice. The difference in the fraction of burst cells at CD4 is not significant in panel **D** after normalization; however, normalization reveals a sex difference in the fraction of active cells at CD4 in panel **B** that was not significant in Fig. 2E. Notably, sex differences at CD55 are no longer significant after normalization. Thus, we conclude that sex differences at CD4 are robust and appear with or without same-mouse normalization, whereas differences at CD55 are less robust.

All values were normalized to the same-mouse mean baseline value. The analysis was identical to Fig. 2D–G, Where the baseline data was compared to each treatment day using Student's t-test with Bonferroni correction for multiple comparisons.

**(E)** Neuronal mean FR was compared between the baseline and TD45 recording sessions. Following treatment, the Clemastine group showed no significant change, whereas the vehicle group showed a significant decrease (p=0.811 and 0.016 for the Clemastine and vehicle groups, respectively, two-tailed paired Student's $t$-test). All values were normalized to the mean value from the corresponding baseline recording.

**(F)** The average number of seconds that fluorescence traces recorded from single neurons exceeded 3 z-scores was not significantly different between baseline and TD45 recording sessions for both Clemastine and vehicle groups (p=0.570 and 0.149 for the Clemastine and vehicle groups, respectively, two-tailed paired Student's $t$-test). All values were normalized to the mean value from the corresponding baseline recording.

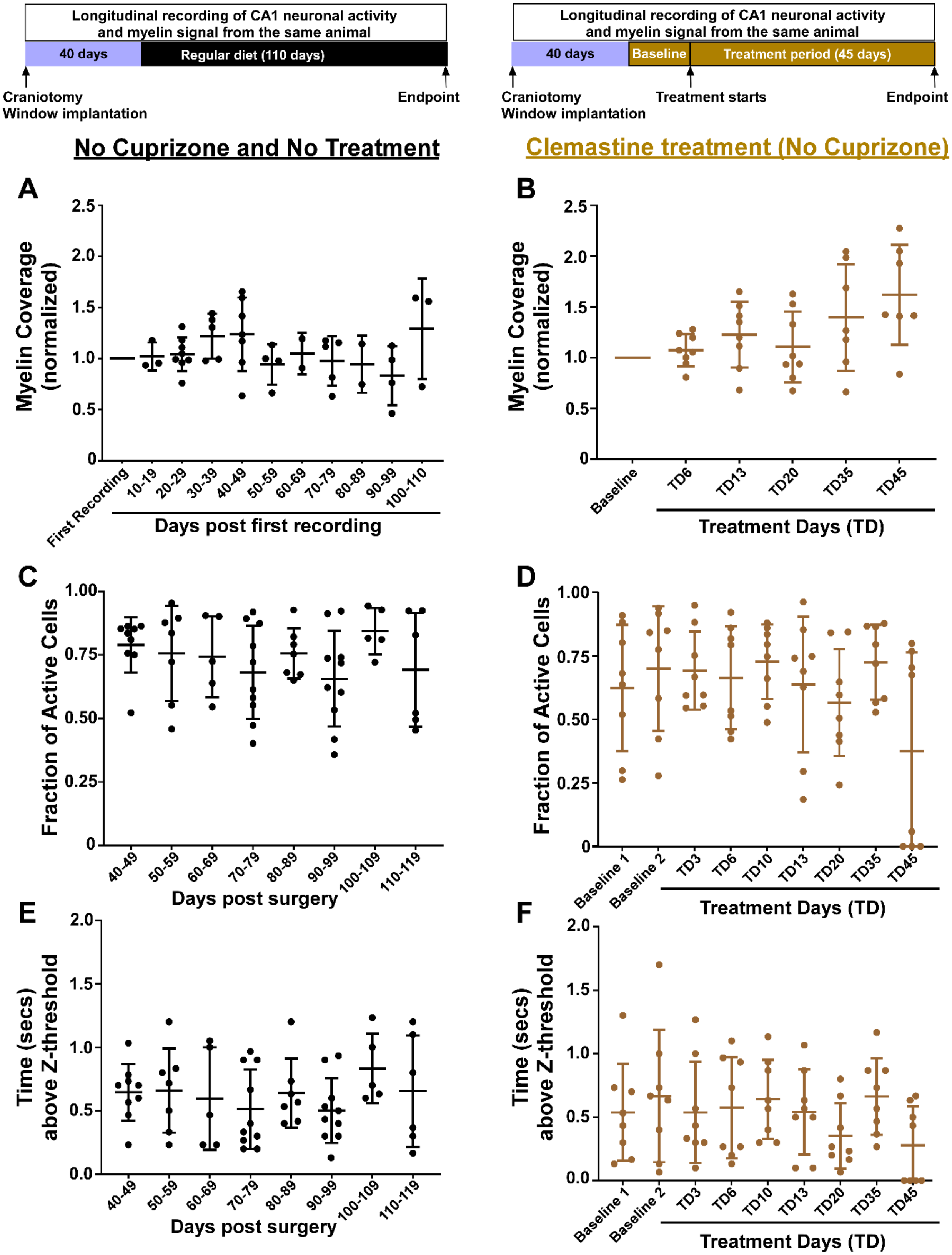
Longitudinal recording of CA1 neuronal activity and myelin signal from the same animal
40 days
Regular diet (110 days)
Craniotomy
Window implantation
Endpoint
Longitudinal recording of CA1 neuronal activity and myelin signal from the same animal
40 days
Baseline
Treatment period (45 days)
Craniotomy
Window implantation
Treatment starts
Endpoint
No Cuprizone and No Treatment
Clemastine treatment (No Cuprizone)
A
Myelin Coverage (normalized)
Days post first recording
B
Myelin Coverage (normalized)
Treatment Days (TD)
C
Fraction of Active Cells
Days post surgery
D
Fraction of Active Cells
Treatment Days (TD)
E
Time (secs) above Z-threshold
Days post surgery
F
Time (secs) above Z-threshold
Treatment Days (TD)

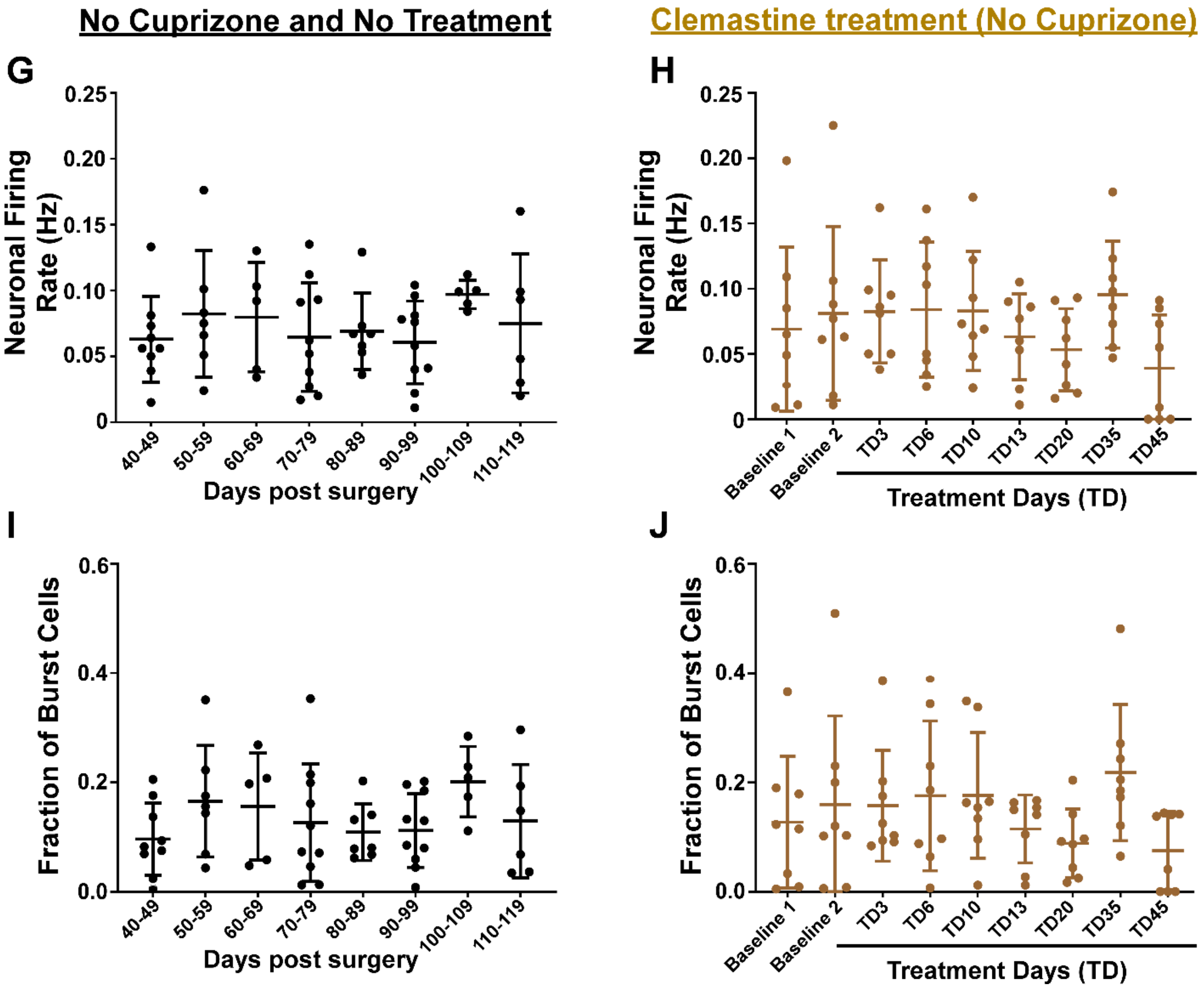


**Figure S3, No significant changes in neuronal firing rate or myelin coverage in control groups, related to Figure 2**

The control cohort included two arms. In the first arm, mice were implanted with hippocampal windows, maintained on a regular diet, and monitored over a time period comparable to the Clemastine- and vehicle-treated mice (n = 10; 7 females, 3 males). For activity recordings, 366–2772 neurons were recorded per mouse (median = 1502), with recording durations of 97–119 days (median = 111 days). For myelin recordings, mice were followed for 39–107 days after the first imaging session (median = 81.5 days). In the second arm, mice were also implanted with hippocampal windows and maintained on a regular diet, but additionally received identical Clemastine treatment for a period matched to that of the experimental mice (n = 8; 5 females, 3 males; recorded at baseline and for 45 days of Clemastine treatment). Monitoring of hippocampal myelination and CA1 activity was identical to the experimental mouse protocol.

**(A-B)** No significant differences in myelination were observed in either control arm. Myelin coverage remained stable in non-treated mice (**A**), whereas a non-significant increase was detected in Clemastine-treated mice (**B**; time effect: $F(9, 26) = 1.998$, $P = 0.07$; mixed-effects model with restricted maximum likelihood). Each dot represents an individual mouse, and bars show mean ± Std. All measurements are normalized to same-mouse baseline values.

**(C-D)** No significant changes were observed in the fraction of active cells in either control condition.

**(E-F)** No significant changes were found in the average number of seconds per recording session during which the neuronal fluorescence signal exceeded 3 z-scores, across all recorded neurons in each mouse, for either control condition.

**(G-H)** No significant changes were observed in mean firing rates during either longitudinal recording or Clemastine treatment in either control condition.

**(I-J)** No significant changes were observed in the fraction of neurons exhibiting burst firing (more than 10 action potentials within a 1-sec time window) in either control condition.

In **A-J**, bold lines represent the mean ± Std, and dots show individual data points.

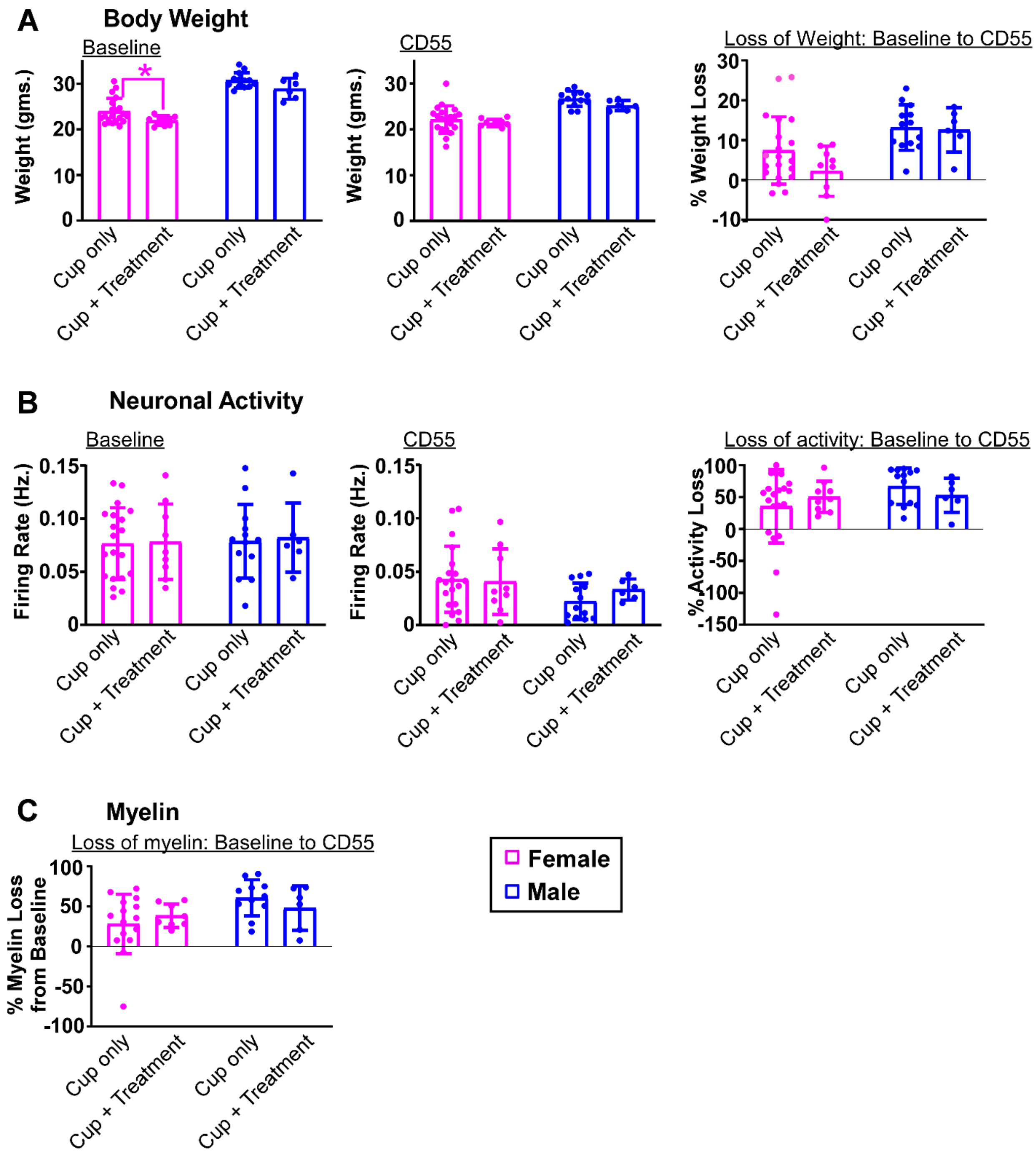


**Figure S4, No relevant differences in body weight, neuronal activity, or myelin status were observed either before or after cuprizone treatment between animals later assigned to Clemastine treatment and those receiving cuprizone alone**

**(A)** ) Body weight at baseline and CD55, and the percent change in body weight at CD55 relative to baseline.

**(B)** Neuronal activity at baseline and CD55, including the percent change in firing rate at CD55 relative to baseline.
**(C)** Percent myelin loss at CD55 relative to baseline.

Comparisons were made between all mice that completed 55 days of cuprizone diet (n = 33; 13 males, 20 females) and all mice that completed 55 days of cuprizone diet followed by 45 days of Clemastine treatment (n = 15; 6 males, 9 females). Each dot represents an individual mouse, and bars show mean ± Std.

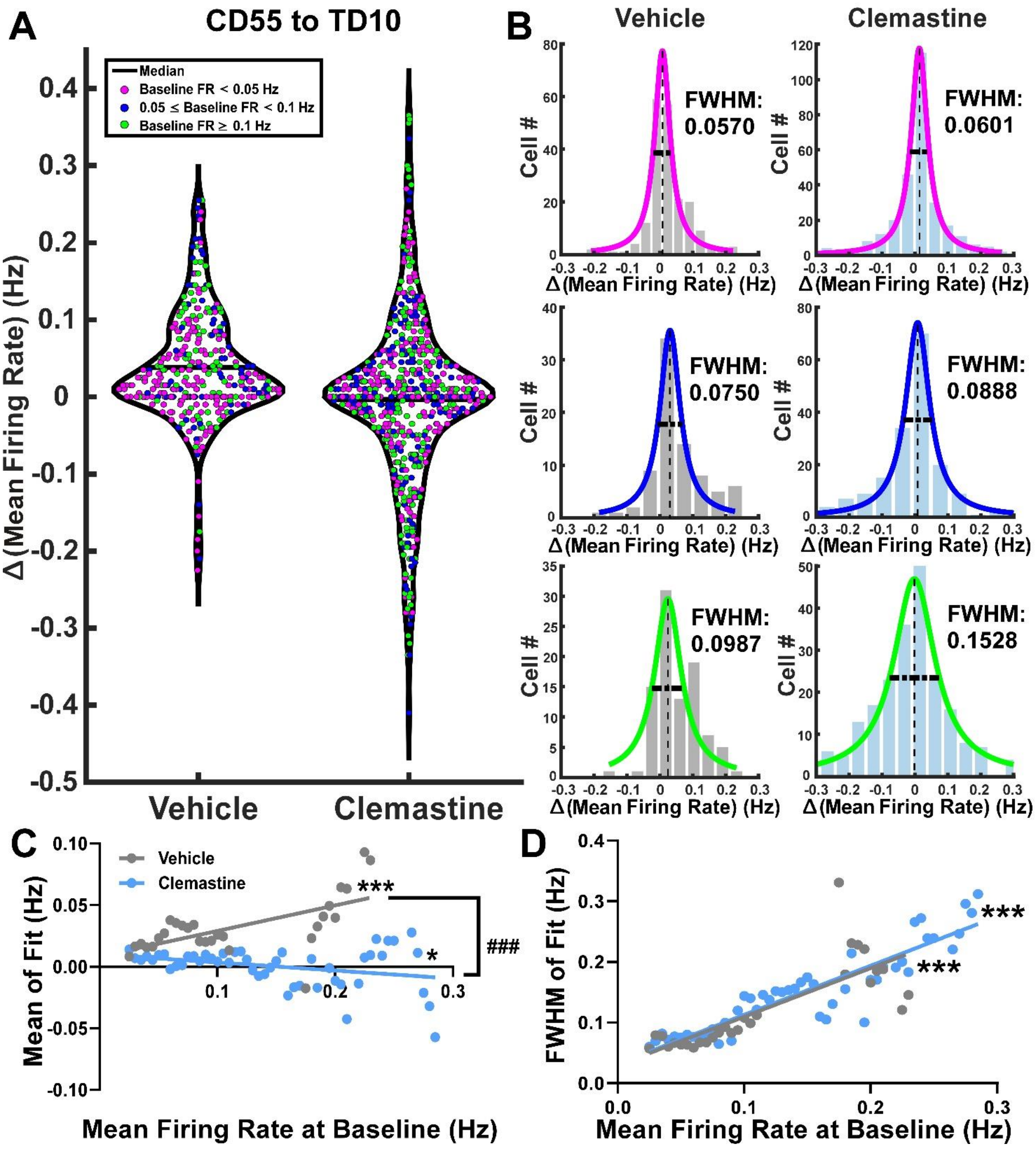


**Figure S5,Single-cell correlates of recovery during the early Clemastine treatment phase, related to Figure 5**

**(A)** Violin plot of the change in mean firing rate of each cell from CD55 to TD10, with each dot colored according to whether the cell's baseline mean firing rate was low, medium, or high (n = 351 and 644 cells from vehicle and Clemastine groups, respectively).

**(B)** Histograms of the data in **A** for low-, medium-, and high-baseline firing-rate cells. Each histogram was fitted with a Lorentzian function as in Fig. 5D, and the mean and FWHM values for each fit were used to assess differences between cell groups. High-activity neurons showed larger FWHM values than medium-activity neurons, which in turn showed larger FWHM values than low-activity neurons (similar to Fig. 5D). However, whereas the mean change in FR for high-activity neurons was positive in the vehicle group, no net change was observed in the Clemastine group.

**(C)** The same cells from **A** were grouped into partially overlapping bins based on their baseline mean FR. For each bin, the change in mean FR from CD55 to TD10 was plotted as a histogram and fitted with a Lorentzian function as in **B**. In vehicle-treated mice, increased activity during the early treatment phase was positively correlated with baseline FR (Pearson's $r = 0.62$, $P < 0.001$). By contrast, Clemastine-treated cells showed a small decrease from CD55 to TD10 that was only weakly related to baseline FR (Pearson's $r = -0.3095$, $P = 0.03$). The slopes of the two regression lines differed significantly (ANCOVA, $P < 0.001$).

**(D)** Both groups showed increased variability (FWHM), which scaled with baseline firing rate (vehicle: Pearson's $r = 0.7903$; Clemastine: $r = 0.9125$; $P < 0.001$ for both groups). The slopes of the two regression lines did not differ significantly.

***, $P < 0.001$, denotes significance of the Pearson correlation; ###, $P < 0.001$, denotes significance of the ANCOVA test for differences in regression slopes.

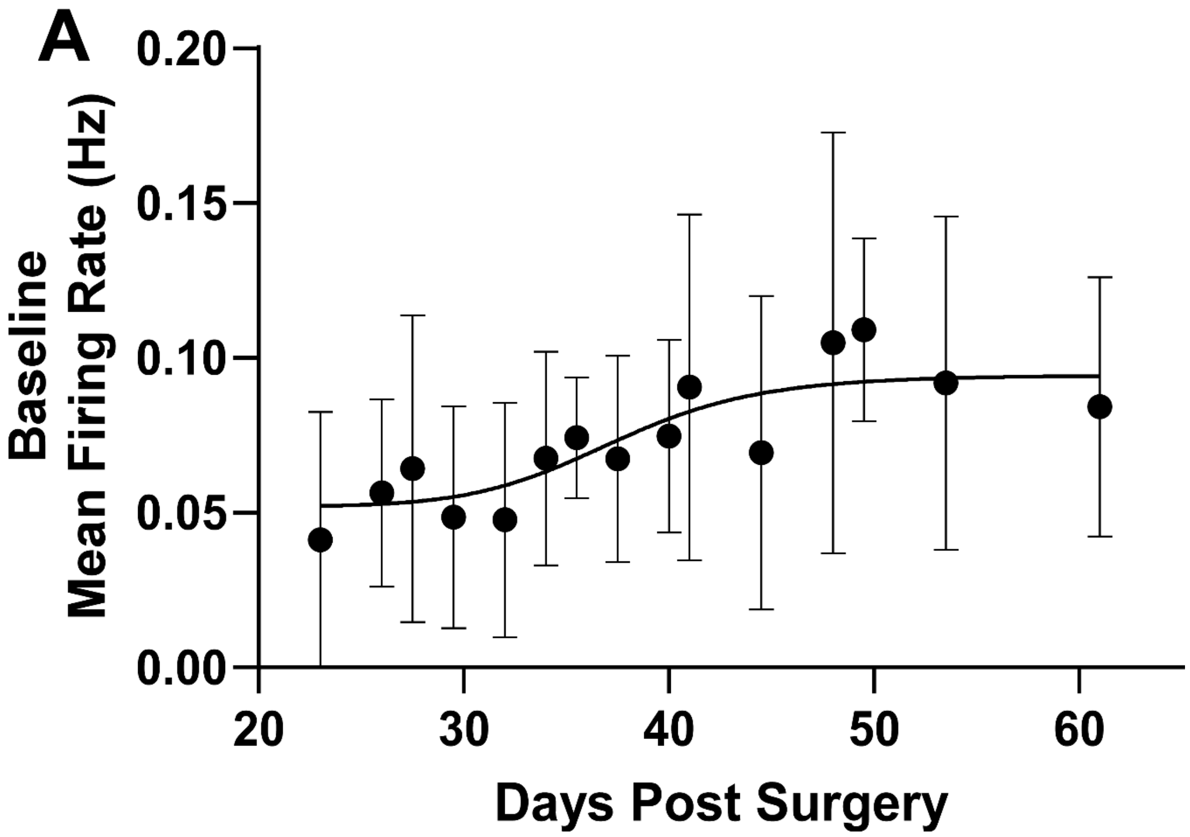


**Figure S6, Baseline firing rate after hippocampal window implantation surgery** Mean FRs from all baseline recordings in all mice (n = 49; 1–5 recordings/mouse; 208–2693 neurons/mouse, median = 1016) were calculated and grouped by the number of days after surgery. Data were binned into 15 temporal bins (7–13 recordings/bin), and the distribution of mean FR in each bin is shown as mean ± Std. A sigmoidal regression line was fit to the data. Based on these findings, a recovery period of at least 40 days after craniotomy was chosen, and mice without at least one baseline recording acquired ≥40 days after surgery were excluded from further analysis (n = 25; 16 males, 9 females).

### Supplementary Statistical Data

| Figure 2 | | F-test statistics | | | P-value | | |
|---|---|---|---|---|---|---|---|
| **Panel** | **Test name** | Time | Group | Time x Group | Time | Group | Time x Group |
| **A** | **Mixed-effects model (REML)** | F (1.876, 66.91) = 14.53 | F (1, 41) = 2.629 | F (1.876, 66.91) = 0.574 | **<0.0001** | **0.1126** | **0.5553** |
| **B** | **Mixed-effects model (REML)** | F (1.911, 72.61) = 30.07 | F (2, 38) = 0.3495 | F (3.822, 72.61) = 12.57 | **<0.0001** | **0.7073** | **<0.0001** |
| **C** | **Mixed-effects model (REML)** | F (1.614, 61.33) = 16.16 | F (2, 38) = 0.5234 | F (3.228, 61.33) = 1.852 | **<0.0001** | **0.5967** | **0.1432** |
| **D** | **Mixed-effects model (REML)** | F (4.556, 207.8) = 19.89 | F (1, 46) = 0.4655 | F (4.556, 207.8) = 1.817 | **<0.0001** | **0.4985** | **0.1177** |
| **E** | **Mixed-effects model (REML)** | F (4.229, 192.9) = 11.85 | F (1, 46) = 1.079 | F (4.229, 192.9) = 2.656 | **<0.0001** | **0.3043** | **0.0315** |
| **F** | **Mixed-effects model (REML)** | F (4.511, 205.7) = 25.62 | F (1, 46) = 0.04805 | F (4.511, 205.7) = 3.703 | **<0.0001** | **0.8275** | **0.0044** |
| **G** | **Mixed-effects model (REML)** | F (3.103, 141.5) = 8.269 | F (1, 46) = 0.04950 | F (3.103, 141.5) = 4.178 | **<0.0001** | **0.8249** | **0.0066** |

| Figure 4 | | F-test statistics | | | P-value | | |
|---|---|---|---|---|---|---|---|
| **Panel** | **Test name** | Time | Group | Time x Group | Time | Group | Time x Group |
| **B** | **Mixed-effects model (REML)** | F (2.188, 36.65) = 21.25 | F (1, 17) = 3.147 | F (2.188, 36.65) = 3.679 | **<0.0001** | **0.094** | **0.0315** |
| **C** | **Mixed-effects model (REML)** | F (3.279, 54.93) = 3.423 | F (1, 17) = 2.476 | F (3.279, 54.93) = 1.177 | **0.0203** | **0.134** | **0.3284** |
| **D** | **Mixed-effects model (REML)** | F (1.504, 9.024) = 0.6414 | F (1, 6) = 0.5011 | F (1.504,9.024) = 3.014 | **0.5063** | **0.5056** | **0.1076** |
| **G** | **Mixed-effects model (REML)** | F (2.728, 32.06) = 0.2268 | F (1, 13) = 1.7 | F (2.728,32.06) = 3.588 | **0.8602** | **0.2149** | **0.0274** |

| Figure S2 | | F-test statistics | | | P-value | | |
|---|---|---|---|---|---|---|---|
| **Panel** | **Test name** | Time | Group | Time x Group | Time | Group | Time x Group |
| **A** | **Mixed-effects model (REML)** | F (2.877, 130.9) = 2.036 | F (1, 46) = 0.0636 | F (2.877, 130.9) = 1.629 | **0.1147** | **0.802** | **0.1876** |
| **B** | **Mixed-effects model (REML)** | F (3.551, 161.6) = 5.082 | F (1, 46) = 0.01077 | F (3.551, 161.6) = 2.609 | **0.0012** | **0.917** | **0.044** |
| **C** | **Mixed-effects model (REML)** | F (3.467, 157.7) = 4.013 | F (1, 46) = 0.243 | F (3.467, 157.7) = 2.613 | **0.006** | **0.6244** | **0.0451** |
| **D** | **Mixed-effects model (REML)** | F (2.144, 97.56) = 0.758 | F (1, 46) = 0.7006 | F (2.144, 97.56) = 1.19 | **0.4798** | **0.4069** | **0.3106** |

| Figure S3 | | F-test statistics | P-value |
|---|---|---|---|
| **Panel** | **Test name** | Time | Time |
| **A** | **One-Way ANOVA** | F (9, 33) = 1.268 | **0.2905** |
| **B** | **Mixed-effects model (REML)** | F (2.153, 14.00) = 3.085 | **0.0747** |
| **C** | **Mixed-effects model (REML)** | F (1.030,6.178) = 1.542 | **0.2608** |
| **D** | **Mixed-effects model (REML)** | F (2.992, 20.94) = 2.904 | **0.059** |
| **E** | **Mixed-effects model (REML)** | F (0.497,2.986) = 1.267 | **0.2736** |
| **F** | **Mixed-effects model (REML)** | F (2.529, 17.71) = 1.655 | **0.2162** |
| **G** | **Mixed-effects model (REML)** | F (2.067,12.40) = 1.094 | **0.367** |
| **H** | **Mixed-effects model (REML)** | F (2.312, 16.18) = 2.175 | **0.1406** |
| **I** | **Mixed-effects model (REML)** | F (2.138,12.83) = 1.231 | **0.3269** |
| **J** | **Mixed-effects model (REML)** | F (2.366, 16.56) = 2.113 | **0.1466** |

| Figure 5 | | R-value | | P-value | |
|---|---|---|---|---|---|
| **Panel** | **Test name** | Vehicle | Clemastine | Vehicle | Clemastine |
| **E** | **Pearson's Rank Correlation** | **-0.8451** | **0.7024** | **<0.001** | **<0.001** |
| **E** | **ANCOVA** | | | **<0.001** | |
| **F** | **Pearson's Rank Correlation** | **0.6602** | **0.8436** | **<0.001** | **<0.001** |
| **F** | **ANCOVA** | | | **<0.001** | |

| Figure S5 | | R-value | | P-value | |
|---|---|---|---|---|---|
| **Panel** | **Test name** | Vehicle | Clemastine | Vehicle | Clemastine |
| **C** | **Pearson's Rank Correlation** | **0.62** | **-0.3905** | **<0.001** | **0.03** |
| **C** | **ANCOVA** | | | **<0.001** | |
| **D** | **Pearson's Rank Correlation** | **0.7903** | **0.9125** | **<0.001** | **<0.001** |
| **D** | **ANCOVA** | | | **n.s.** | |

**Table S1.** Statistical values for linear mixed-effects, correlation, and ANCOVA tests.

**Figure 2- Post-hoc test: Multiple Comparisons**

| Panel | Comparisons | Test | P Value |
|---|---|---|---|
| A | BL vs CD46 (Female) | Bonferroni | 0.0008 |
| | BL vs CD55 (Female) | Bonferroni | 0.0003 |
| | CD18 vs CD46 (Female) | Tukey | 0.016 |
| | CD18 vs CD55 (Female) | Tukey | 0.0018 |
| | BL vs CD32 (Male) | Bonferroni | 0.02 |
| | BL vs CD46 (Male) | Bonferroni | <0.0001 |
| | BL vs CD55 (Male) | Bonferroni | <0.0001 |
| | CD18 vs CD55 (Male) | Tukey | <0.0001 |
| | CD32 vs CD55 (Male) | Tukey | 0.0002 |
| B | CD55 (Male vs Control) | Tukey | 0.0001 |
| | CD55 (Female vs Control) | Tukey | 0.0002 |
| | Male (BL vs CD32) | Tukey | 0.0287 |
| | Male (BL vs CD55) | Tukey | <0.0001 |
| | Male (CD32 vs CD55) | Tukey | 0.0005 |
| | Female (BL vs CD32) | Tukey | 0.0114 |
| | Female (BL vs CD55) | Tukey | 0.0004 |
| | Female (CD32 vs CD55) | Tukey | 0.0047 |
| C | Male (BL vs CD55) | Tukey | 0.0035 |
| | Female (BL vs CD55) | Tukey | 0.0108 |
| | Female (CD32 vs CD55) | Tukey | 0.0182 |
| D | BL vs CD4 (Female) | Tukey | <0.0001 |
| | BL vs CD18 (Female) | Tukey | 0.0004 |
| | BL vs CD32 (Female) | Tukey | 0.0178 |
| | BL vs CD46 (Female) | Tukey | 0.0005 |
| | BL vs CD55 (Female) | Tukey | 0.0002 |
| | CD4 vs CD32 (Female) | Tukey | 0.0111 |
| | CD4 vs CD46 (Female) | Tukey | 0.0189 |
| | BL vs CD4 (Male) | Tukey | 0.0114 |
| | BL vs CD18 (Male) | Tukey | 0.0035 |
| | BL vs CD32 (Female) | Tukey | 0.0293 |
| | BL vs CD46 (Male) | Tukey | 0.0032 |
| | BL vs CD55 (Male) | Tukey | 0.0005 |

**Figure 2- Post-hoc test: Multiple Comparisons**

| Panel | Comparisons | Test | P-value |
|---|---|---|---|
| E | BL vs CD4 (Female) | Tukey | <0.0001 |
| | BL vs CD18 (Female) | Tukey | 0.0131 |
| | BL vs CD46 (Female) | Tukey | 0.0244 |
| | BL vs CD55 (Female) | Tukey | 0.0036 |
| | CD4 vs CD32 (Female) | Tukey | 0.0021 |
| | BL vs CD46 (Male) | Tukey | 0.012 |
| | BL vs CD55 (Male) | Tukey | 0.0021 |
| F | CD4 (Male vs Female) | Tukey | 0.0036 |
| | CD55 (Male vs Female) | Tukey | 0.0185 |
| | BL vs CD4 (Female) | Tukey | <0.0001 |
| | BL vs CD18 (Female) | Tukey | <0.0001 |
| | BL vs CD32 (Female) | Tukey | 0.0055 |
| | BL vs CD46 (Female) | Tukey | 0.0001 |
| | BL vs CD55 (Female) | Tukey | <0.0001 |
| | CD4 vs CD32 (Female) | Tukey | 0.0084 |
| | BL vs CD4 (Male) | Tukey | 0.0014 |
| | BL vs CD18 (Male) | Tukey | 0.0011 |
| | BL vs CD32 (Female) | Tukey | 0.0074 |
| | BL vs CD46 (Male) | Tukey | 0.0008 |
| | BL vs CD55 (Male) | Tukey | <0.0001 |
| | CD4 vs CD55 (Male) | Tukey | 0.0074 |
| | CD32 vs CD55 (Male) | Tukey | 0.0339 |
| G | CD4 (Male vs Female) | Tukey | 0.0144 |
| | CD55 (Male vs Female) | Tukey | 0.0439 |
| | BL vs CD4 (Female) | Tukey | 0.0004 |
| | BL vs CD18 (Female) | Tukey | 0.0037 |
| | BL vs CD32 (Female) | Tukey | 0.0048 |
| | BL vs CD46 (Female) | Tukey | 0.0007 |
| | BL vs CD18 (Male) | Tukey | 0.0349 |
| | BL vs CD32 (Female) | Tukey | 0.0091 |
| | BL vs CD46 (Male) | Tukey | 0.0136 |
| | BL vs CD55 (Male) | Tukey | 0.0013 |
| | CD4 vs CD55 (Male) | Tukey | 0.0053 |
| | CD18 vs CD55 (Male) | Tukey | 0.0477 |

**Figure 4- Post-hoc test: Multiple Comparisons**

| | | | |
|---|---|---|---|
| B | TD13 (Clemastine vs Vehicle) | Tukey | 0.0291 |
| | Clemastine (BL vs CD55) | Tukey | 0.0014 |
| | Clemastine (BL vs TD6) | Tukey | 0.003 |
| | Clemastine (BL vs TD13) | Tukey | 0.0005 |
| | Clemastine (CD55 vs TD45) | Tukey | 0.0476 |
| | Vehicle (BL vs CD55) | Tukey | 0.0311 |
| | Vehicle (BL vs TD6) | Tukey | 0.03 |
| | Vehicle (BL vs TD45) | Tukey | 0.0383 |
| G | TD45 (Clemastine vs Vehicle) | Tukey | 0.0495 |

**Figure S2- Post-hoc test: Multiple Comparisons**

| Panel | Comparisons | Test | P-value |
|---|---|---|---|
| A | BL vs CD4 (Female) | Bonferroni | <0.0001 |
| | BL vs CD55 (Female) | Bonferroni | 0.008 |
| | BL vs CD18 (Male) | Bonferroni | 0.003 |
| | BL vs CD46 (Male) | Bonferroni | 0.043 |
| | BL vs CD55 (Male) | Bonferroni | 0.001 |
| B | CD4 (Male vs Female) | Tukey | 0.0346 |
| | BL vs CD4 (Female) | Bonferroni | <0.0001 |
| | BL vs CD55 (Female) | Bonferroni | 0.007 |
| | CD4 vs CD32 (Female) | Tukey | 0.0075 |
| | BL vs CD18 (Male) | Bonferroni | 0.042 |
| | BL vs CD46 (Male) | Bonferroni | 0.0027 |
| | BL vs CD55 (Male) | Bonferroni | 0.0005 |
| | CD32 vs CD55 (Male) | Tukey | 0.0343 |
| C | CD4 (Male vs Female) | Tukey | 0.0243 |
| | BL vs CD4 (Female) | Bonferroni | <0.0001 |
| | BL vs CD18 (Female) | Bonferroni | 0.0011 |
| | BL vs CD46 (Female) | Bonferroni | 0.0016 |
| | BL vs CD55 (Female) | Bonferroni | <0.0001 |
| | BL vs CD4 (Male) | Bonferroni | <0.0001 |
| | BL vs CD18 (Male) | Bonferroni | 0.0017 |
| | BL vs CD46 (Male) | Bonferroni | <0.0001 |
| | BL vs CD55 (Male) | Bonferroni | <0.0001 |
| | CD4 vs CD55 (Male) | Tukey | 0.0049 |
| D | BL vs CD46 (Male) | Bonferroni | 0.001 |
| | BL vs CD55 (Male) | Bonferroni | <0.0001 |

**Table S2:** Statistical test values of post-hoc comparisons.

| Group | Mouse ID | Exclusion decision made on |
|---|---|---|
| **Cuprizone** | A1230 | Baseline |
| | A1235 | CD18 |
| | A1251 | CD18 |
| | A1309 | CD4 |
| | A1310 | CD4 |
| | A1320 | CD4 |
| | A1321 | CD4 |
| | A1336 | CD4 |
| | A1337 | CD46 |
| | A1342 | CD46 |
| | A1344 | CD46 |
| | A1345 | CD46 |
| | A1346 | CD46 |
| | A1349 | CD18 |
| | A1350 | Baseline |
| | A1372 | CD4 |
| | A1376 | CD4 |
| | A1388 | Baseline |
| | A1395 | Baseline |
| | A1436 | CD18 |
| | A1468 | Baseline |
| | A1463 | Baseline |
| | A1244 | CD32 |
| **Clemastine** | A1252 | TD24 |
| | A1335b | TD13 |
| | A1438 | TD10 |
| **Vehicle** | A1239 | TD6 |
| | A1241 | TD3 |
| | A1274 | TD6 |
| | A1341 | TD17 |
| | A1432 | TD10 |
| | A1461 | TD10 |
| | A1464 | TD3 |
| **Control** | A1340 | Day 46 |
| | A1417 | Day 71 |
| | A1445 | Day 49 |

**Table S3: List of excluded mice based on their health and window condition status.** The table lists the mice that were completely removed from the study because they didn't complete the cuprizone or treatment phase in their respective group, as well as the recording date where the decision was taken (such as first signs of health concerns or loss of the cranial window). For mice in the control group, the days were counted from the date of craniotomy.

| Group | Mouse ID | Excluded recording days |
|---|---|---|
| **Cupriozne** | A1226 | CD46 (TPLSM and THG) |
| | A1232 | CD 55 (THG) |
| | A1233 | CD46 (THG) |
| | A1239 | CD46 (THG) |
| | A1333 | CD18 (THG) |
| | A1335b | CD46, CD55 (THG) |
| | A1371 | CD18 (THG) |
| | A1435 | CD18 (THG) |
| | A1438 | CD46, CD55 (THG) |
| | A1461 | CD46 (THG) |
| | A1464 | CD32, CD46 (THG) |
| **Clemastine** | A1224 | CD46 (TPLSM) |
| | A1242 | TD25 (TPLSM) |
| | A1465 | CD46 (THG) and TD17 (TPLSM) |
| | A1469 | CD46 (THG) and TD17 (TPLSM) |
| **Vehicle** | A1312 | TD13 (TPLSM) |
| | A1313 | TD13 (TPLSM) |
| **Control (No Cup.)** | A1363 | Day 156 (THG) |
| | A1403 | Days 113 (TPLSM), 140 (THG) |
| | A1419 | Days 129 (TPLSM), 156 (THG) |
| | A1448 | Days 174, 188 (THG) |
| | A1453 | Day 93 (THG) |
| **Control (Clem. Only)** | A1518 | TD35, TD45 (THG) |

**Table S4: Exclusion of specific recording sessions due to quality issues.** The table lists mice and specific recording sessions that were excluded from the analysis due to quality issues. Quality assessment was independently conducted for neuronal activity recording (TPLSM) and myelin monitoring (THG). Additionally, THG recording data from seven mice was entirely excluded due to consistent quality issues. For mice in the control group, the days were counted from the date of craniotomy.
